# dMSCC: A microfluidic platform for microbial single-cell cultivation under dynamic environmental medium conditions

**DOI:** 10.1101/2020.07.10.188938

**Authors:** Sarah Täuber, Corinna Golze, Phuong Ho, Eric von Lieres, Alexander Grünberger

## Abstract

In nature and in technical systems, microbial cells are often exposed to rapidly fluctuating environmental conditions. These conditions can vary in quality, e.g., existence of a starvation zone, and quantity, e.g., average residence time in this zone. For strain development and process design, cellular response to such fluctuations needs to be systematically analysed. However, the existing methods for physically emulating rapidly changing environmental conditions are limited in spatio-temporal resolution. Hence, we present a novel microfluidic system for cultivation of single cells and small cell clusters under dynamic environmental conditions (dynamic microfluidic single-cell cultivation (dMSCC)). This system enables to control nutrient availability and composition between two media with second to minute resolution. We validate our technology using the industrially relevant model organism *Corynebacterium glutamicum*. The organism was exposed to different oscillation frequencies between nutrient excess (feasts) and scarcity (famine). Resulting changes in cellular physiology, such as the colony growth rate and cell morphology were analysed and revealed significant differences with growth rate and cell length between the different conditions. dMSCC also allows to apply defined but randomly changing nutrient conditions, which is important for reproducing more complex conditions from natural habitats and large-scale bioreactors. The presented system lays the foundation for the cultivation of cells under complex changing environmental conditions.

## Introduction

In large scale industrial cultivations, but also in the natural habitat, microbial cells are exposed to rapidly changing conditions in quality, e.g., existence of a starvation zone, and quantity, e.g., average residence time in this zone.^1–3^ These conditions can vary in temperature, oxygen, pH value, shear rate etc. The fluctuating environmental conditions in bioreactors can be caused by long mixing times due to the size of the reactor, the geometry of the reactor (stirrer, feeding, gassing) and the operating mode (speed, feeding, gassing). This creates gradients of pH value and nutrients, e.g. substrate concentration and dissolved oxygen.^1^ In bioreactors, highly concentrated nutrients are fed regularly, creating different zones of nutrients in the reactor, e.g. zones with substrate excess and zones where no substrates are available. As a result, cells are exposed to different environmental conditions over time, which can be represented by lifelines (recording the environment over time).^4^ Typical time scales are in second to minutes range based on the size and geometry of the bioreactor.^5^ This has an impact on various parameters, which can be divided into biological and technical variables.^6^ Biological parameters include the influence on cellular metabolism on different cellular levels (transcription, translation)^7,8^, while technical parameters include for example productivity and yield.^9^

The analysis of cellular behaviour upon defined environmental perturbation on single-cell level is of central interest for the understanding of fundamental topics such as growth rates^10,11^, population stability^12^ and cellular adaptation^13^. Furthermore, biotechnological questions on the behaviour of single cells in the bioprocess are decisive, e.g., how cells respond to gradients in bioreactors and how cells adapt their gene network upon environmental changes. In order to understand these phenomena, novel methods must be developed, that allow the cultivation of cells at dynamic environmental conditions at the single-cell level.

In recent years, many microfluidic devices have been developed and applied for the cultivation and analysis at single-cell level under defined environmental conditions.^14–16^ Here, cells are trapped in different cultivation chamber geometries and perfused with medium. The cultivation chambers on these microfluidic devices range from 0D to 3D systems.^14^ For most of the reported single-cell cultivation studies 1D and 2D designs were used. In 1D systems cell growth is enabled in a line of cells upon 4 – 10 cells which allows long-term cultivations of small sub-populations. 2D systems allow to cultivate small cell colonies up to 1000 cells in a monolayer.

New microfluidic systems are currently being developed for the analysis of cellular behaviour under dynamic environmental conditions.^17,18^ The technical setups (pumps and valves) for the generation of dynamic environmental conditions can be implemented outside the microfluidic device (e.g. pressure driven pumps) or integrated into the microfluidic device (e.g. pneumatic valves).^17^ Various environmental profiles can be created in these microfluidic systems with different modulation modes. First, the frequency of the pulses can be varied which is called pulse frequency modulation (PFM). Here, the width between the pulses and the amplitude of the pulses are constant. Second, the width of the pulses can be varied, i.e. the time when no new pulse is inserted (pulse width modulation (PWM)). Here, the frequency of the pulses and the amplitude are constant. Third, the amplitude of the pulses can be changed (pulse amplitude modulation (PAM)). Furthermore, random profiles can be generated.

In the last years, several microfluidic approaches for studies of cellular physiology at dynamic environmental conditions have been reported (Table 1). Kaiser et al.^19^ developed a dial-a-wave system (combination of hardware, software and microfluidic device which include a fluidic mixer) and cultivated *E. coli* cells in mother machine growth channels (1D) with a periodic oscillation of glucose and lactose in 4 hour intervals (PFM-mode). They could show that cells expressed lacZ-GFP from the lac promotor by switching the carbon source from glucose to lactose. Cells completely stopped their growth after three minutes when a switch from glucose to lactose was performed. Rojas et al.^20^ used the commercially available microfluidic CellASIC system for the cultivation of *E. coli* cells with a periodic oscillation of LB+sorbitol and LB medium in 45 second intervals (PFM-mode). They analysed single-cell behaviour of *E. coli* cells upon hyperosmotic shocks and found that cell elongation is slowed down. When the turgor is restored, the length of the cells recovers. Lugagne et al.^21^ cultivated *E. coli* in 1D growth channels with 3.5 h oscillation time including 3 h intervals of LB medium + IPTG and 30 min intervals of LB medium + aTc (PWM-mode). They analysed the gene regulation in a bistable genetic toggle switch in *E. coli* and showed that this can be maintained dynamically. Lambert et al.^22^ also cultivated *E. coli* in 1D growth channels with an oscillation between MOPS minimal medium + 0.4% lactose and MOPS minimal medium + 0.4% glucose in 5 minute to 4 hour intervals (PFM-mode). They analysed the non-genetic memory in *E. coli* and showed that the phenotype memory reduced the lag phase under oscillating environment.

**Table 1:** Overview of different environmental profiles and conditions of microfluidic single-cell cultivation with microbial cells.

| Environmental profile (period) | Environmental conditions | Growth chamber design | Application | Ref. |
| --- | --- | --- | --- | --- |
| PFM (4 h; 4h) | Glucose; lactose | 1D | Analysis of lac operon response in *E. coli* | <sup>19</sup> |
| PFM (45 sec; 45 sec) | LB; LB medium + sorbitol | 2D | Investigation of hyperosmotic shock in *E. coli* | <sup>20</sup> |
| PWM (3h; 30 min) | LB medium + IPTG; LB medium + aTc | 1D | Investigation of gene regulation in *E. coli* (bistable genetic toggle) | <sup>21</sup> |
| PFM (5 min; 5 min – 4h;4h) | MOPS minimal medium + 0.4% lactose; MOPS minimal medium + 0.4% glucose | 1D | Investigation of non-genetic memory in *E. coli* | <sup>22</sup> |
| PFM + PWM (10sec;10sec – 1h;1h) | BHI medium; PBS buffer | 2D | Analysis of growth behaviour in *C. glutamicum* | This work |

Most of the demonstrated examples were developed and applied for the investigation of cellular physiology using PFM with frequencies between 5 min and hours. Studies with oscillations in seconds to minutes range have not been performed systematically yet. Recently, Ho et al.^23^ showed that the cultivation of cells with a combination of PWM, PFM and PAM with second scale resolution is challenging, depending on the cultivation chamber design and applied periphery. Thus, systems need to be carefully designed and the feasibility of fast environmental changes needs to be shown for each setup and configuration to guarantee defined medium oscillations.

In this work, we present a microfluidic single-cell cultivation system for the cultivation of microbial cells under dynamic environmental conditions (dynamic microfluidic single-cell cultivation (dMSCC)). Due to the design of the microfluidic system, cells can be cultivated with a high degree of parallelization allowing the collection of statistically meaningful information. The dMSCC setup can create oscillating nutrient profiles between two specified medium conditions, i.e. amount and composition, with a high frequency and thus enables a wide range of experiments with PFM and PWM. Moreover, control measurements of the different medium conditions can be performed. Experimental as well as computational methods were used to characterize mass transfer during the dMSCC and investigate the feasibility of realizing controlled, oscillating conditions in the second range. The cultivation of *Corynebacterium glutamicum* with different oscillation frequencies between nutrient excess (complex BHI medium) and scarcity (PBS buffer) were performed as a model study.

Furthermore, a first proof of concept for the realization of complex environmental profiles as present in bioreactors was demonstrated.

## Material and methods

### Pre-cultivation, bacterial strain and growth media

The bacterial strain *C. glutamicum* strain WT ATCC 13032 was used in this study. *C. glutamicum* was cultivated in BHI (Brain heart infusion, Roth, Germany) medium using 37 g L^−1^ BHI at 30 °C. The medium was autoclaved and additionally sterile filtered to prevent channel clogging during the microfluidic experiments.

Overnight pre-cultures of *C. glutamicum* were inoculated from glycerol stock in 10 mL BHI medium in 100 mL flasks on a rotary shaker at 120 rpm. Cells from the overnight culture were transferred to inoculate the second culture with a starting OD_600_ of 0.05. When the culture had an OD_600_ of around 0.2 the cells were seeded in the microfluidic device. After seeding, the cells were dynamically perfused with BHI medium and PBS buffer (8 g L^−1^ NaCl, 0.2 g L^−1^ KCl, 1.42 g L^−1^ Na_2_HPO_4_, 0.27 g L^−1^ KH_2_PO_4_) according to the protocol description below.

### Wafer fabrication

A silicon wafer mould was fabricated with a two-layer photolithography process and served as a mould for PDMS soft lithography.^24^ In the following the major processing steps and parameters are described.

Photolithography was performed under cleanroom condition. A 4-inch silicon wafer was cleaned with permonosulphuric acid and demineralised water. Afterwards, the wafer was spin coated and dehydrated through a 15-minute dehydration bake at 200 °C. A 800 nm thick layer of negative photoresist SU-8 (18% solid) was spin coated and was pre-bake at 65 °C for 1 minute. Photolithography was performed with a laser beam written 4-inch lithography mask (Deltamask, Netherlands). The exposure time was optimized for the SU-8 thickness, structure resolution and lamp intensity of the mask aligner (MJB3, Süss MicroTec, Germany). Exposure time was set to 1.3 seconds in vacuum contact mode. Furthermore, a 5-minute post bake at 65 °C and 95 °C was performed and the wafer was developed in a negative resist developer (mrdev 600, micro resist technology GmbH, Germany). Afterwards, a second SU-8 layer (52% solid) with a thickness of 10 µm was spin coated onto the wafer. The lithography process was equal to the first layer except the pre-bake process, which was set to 5 minutes at 65 °C and 5 minutes at 95 °C and the exposure time, which was adjusted to 6 seconds.

### PDMS Moulding

The wafer was covered with PDMS in a ratio 10:1 between base and curing agent (Sylgard 184 Silicone Elastomer, Dow Corning Corporation, USA). Afterwards, the wafer was degassed in an exicator for 30 minutes and backed at 80 °C for 2 hours (universal cupboard, Memmert GmbH, Germany). After this step, the PDMS chips were cut out from the wafer and cleaned three times with isopropanol and blown dry with pressurized air. The cover glasses (D 263 T eco, 39.5×34.5×0.175 mm, Schott, Germany) for the microfluidic chip were also cleaned in this step. Afterwards, the PDMS chip and the cover glass were oxygenised with O_2_ plasma (Femto Plasma Cleaner, Diener Electronics, Ebhausen, Germany) for 24 seconds with a power of 45% and assembled. Prior the use PDMS-glass bonding was strengthened by a 2-minute bake at 80 °C.

### Generation of oscillation profiles

#### Oscillation setup

The microfluidic device was flushed with a cell suspension which was transferred from the main culture at exponential phase with maximum OD_600_ of 0.2. Flow was stopped when enough cultivation chambers were seeded with a single cell. Then, the flow was switched from bacterial suspension to growth medium and buffer. Medium flow was achieved with high precision pressure pumps (Line-up series, Fluigent, Jena, Germany) with a pressure of 150 and 75 mbar (flow rate of ∼ 6.2 µl/ min). Medium was switched with an automated software tool (microfluidic automation tool (MAT), Fluigent, Jena, Germany).

#### Flow characterisation

The flow characterisation within the dMSCC setup was performed with coloured ink (Hardtmuth set of technical inks, Koh-I-Noor, Czech Republic). Different pump settings were tested for the optimal oscillation flow profiles in the microfluidic device. Experimental validation of the mass transfer between supply channels and cultivation chambers was verified with fluorescein (Macrolex yellow, Lanxess, Germany). Medium exchange was characterized by the course of fluorescence upon a change between fluorescein and ethanol. The highest value of the fluorescein signal in a chamber was set to 100% and no fluorescence in the chamber to 0%.

#### CFD simulations

Computational fluid dynamics (CFD) simulations were conducted in COMSOL Multiphysics 5.4.0.225 (COMSOL AB, Sweden). In the CFD model, glucose was used as a representative nutrient, which was specified in the defined CGXII growth medium.^25^ Both in the defined medium and in the complex medium the glucose concentration is supersaturated. The maximum substrate concentration was as low as 222 mol L^−1^. The stationary velocity field and transient mass transport of glucose were calculated in order to track the extracellular substrate concentration in the cultivation chamber. The CFD analysis was performed for one representative, empty chamber (see SI Material and Methods). The velocity and concentration fields were sequentially computed and thus the flow field can be regarded as practically independent of the concentration field. For comparison purposes, the medium exchange was described by the course of glucose concentration in the whole cultivation chamber.

#### Irregular oscillation profiles – lifelines

Lifelines simulated by Haringa et al.^4^ were chosen to be reproduced by dMSCC. Here, PBS buffer was pulsed on BHI medium. The PBS pulses length varied in all profiles in the following order: 60 sec, 5 sec, 25 sec, 15 sec, 50 sec, 10 sec, 45 sec, 30 sec, 60 sec and 5 sec, only the BHI width between theses pulses were varied to generate three different profiles, that differed in the overall medium supply (40% – 80%) during the dMSCC.

#### Time-lapse imaging

Time-lapse microscopy was performed using an inverted automated microscope from Nikon (Nikon Eclipse Ti2, Nikon, Germany). The microscope stage was surrounded with a cage incubator for the optimal temperature control (Cage incubator, OKO Touch, Okolab S.R.L., Italy). The microfluidic device was placed inside the cage incubator in an in-house fabricated chip holder. Additionally, the setup was equipped with an 100x oil objective (CFI P-Apo DM Lambda 100x Oil, Nikon GmbH, Germany), DS-Qi2 camera (Nikon camera DS-Qi2, Nikon GmbH, Germany) and an automated focus system (Nikon PFS, Nikon GmbH, Germany) to compensate the thermal drift during long term microscopy. 60 cultivation chambers were selected manually for each experiment and were managed with NIS-Elements Imaging Software (Nikon NIS Elements AR software package, Nikon GmbH, Germany). Time-lapse images were recorded every 10 minutes.

## Data analysis

Data analysis of the live-cell image sequences was performed by using the open source software Fiji^26^. First, a k-mean clustering for the background correction was performed. In the phase contrast images for each time point the cells were separated from the background. The cluster with the cells was retained, the background and the intermediate area between the cells were removed. The cell contours were flattened by morphological operators such as dilatation. Cells which were too close together could not be separated by clustering but were segmented by watershed transformation. Afterwards, the particle analyser was used to obtain the area and the cell count for each time point.

The growth rate was determined by a linear regression through the semi-logarithmic plot of cell numbers over the time. The slope indicates the growth rate.

## Results and Discussion

### Device principle and design

The presented microfluidic device (Fig. 1A) was designed for the cultivation and analysis of single cells and cell clusters under dynamic environmental conditions. We designed a chip system containing several arrays of monolayer cultivation chambers, further denoted as cultivation chamber, connected to two medium inlet channels. Each microfluidic chip contains three parallel, separate cultivation units.

**Fig. 1:**
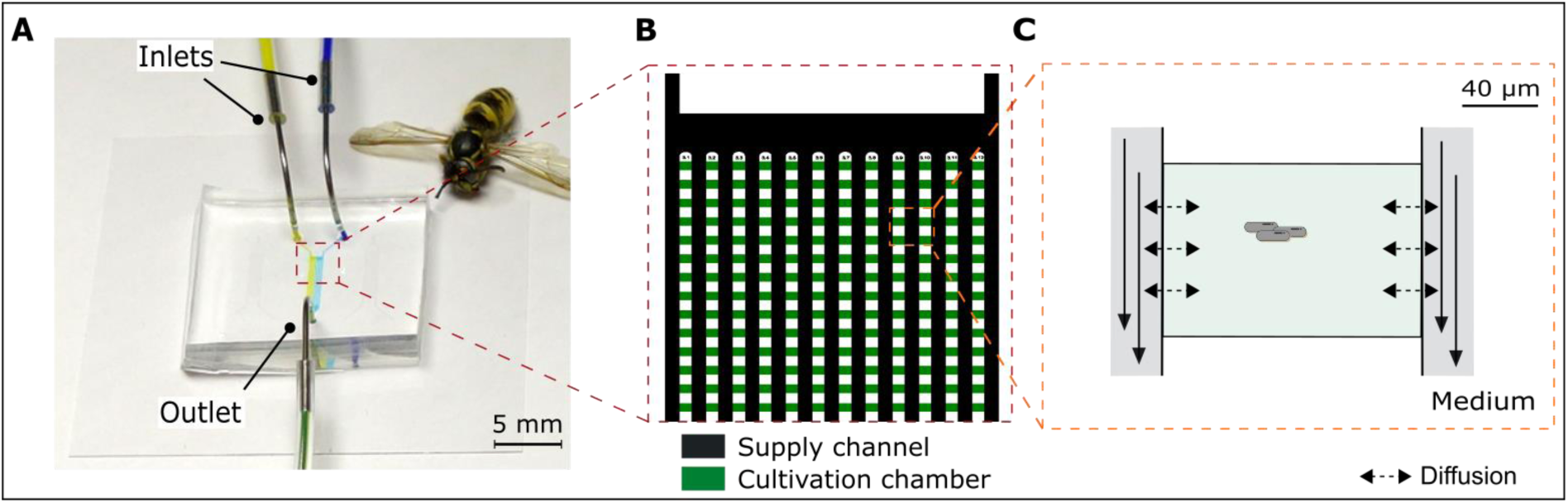
Design of the microfluidic chip for cultivation of single cells and small cell clusters under dynamic environmental conditions. A) Microfluidic chip with two inlets and one outlet per cultivation unit. B) Illustration of the 12 arrays of cultivation chambers. C) Individual monolayer cultivation chamber.

Medium inlet channels are placed at the left and right corner of the 12 cultivation chamber arrays (Fig. 1B). Each cultivation chamber array (a linear series of chambers) has 50 monolayer cultivation chambers. The cultivation chambers are designed as open box chambers (Fig. 1C). The monolayer growth chamber design is based on Grünberger et al.^27^, with cultivation chamber dimensions of 90 µm × 80 µm, covering an area of around 7200 µm^2^ per cultivation chamber. This enables the cultivation of microcolonies up to several hundred of cells. The cultivation chambers (Fig. 1B green) are around 800 nm in height which restricts cell growth to a monolayer. The flow inside the supply channel is laminar and the mass transfer occurs mainly via diffusion inside the cultivation chambers.^28^ The arrays are connected to supply channels (Fig. 1B black). The supply channels have a height of around 10 µm and a width of 100 µm.

### Dynamic environmental conditions

#### Technical Principle

The working principle of our dMSCC system is illustrated in Fig. 2. During dMSCC experiments, medium is switched dynamically between two reservoirs, leading to three distinct cultivation zones (Fig. 2). The two zones are classified as the control zones I & II (left and right) and one zone contains the arrays where medium can be (arbitrarily) changed between the values of the control zones (switching zone). Flow profiles are adjusted in such a way, that always three arrays on the left and right side are control zones and six arrays belong to the switching zone, where cultivation conditions are regulated dynamically. The area where the two liquids converge creates a sharp border layer based on the laminar flow. Fig. 2A shows the flow profile I where nine cultivation chamber arrays are flushed with liquid A and three cultivation chamber arrays as negative control (control zone II) are flushed with liquid B. Fig. 2B shows the flow profile II, where the six cultivation chamber arrays in the centre are perfused with liquid B so that only the positive control (control zone I) is flushed with liquid A. During dMSCC, the system is repeatedly switched between flow profile I and II in specific time-intervals. Fig. 2C and D show the experimental validation of the flow profiles with coloured ink (blue for liquid A and red for liquid B). Furthermore, we tested the spatial reproducibility of the oscillation with a rectangular flow profile between blue and red colour ink in 10 second intervals (see Fig. S1). Here, the deviation (Δx) of the fluid boundaries of the two liquid streams, which are formed by the laminar flow profile, was analysed (see Fig. S1A) and resulted in a deviation from the setpoint Δx = 0 between – 8 µm and 7 µm (see Fig. S1B). The deviation is most likely caused by the external compressed air fluctuations that are applied to the pressurized pumps. Due to the main channel width of 100µm, the detected deviation of ∼ 7 µm can be neglected for the dMSCC experiments.

**Fig. 2:**
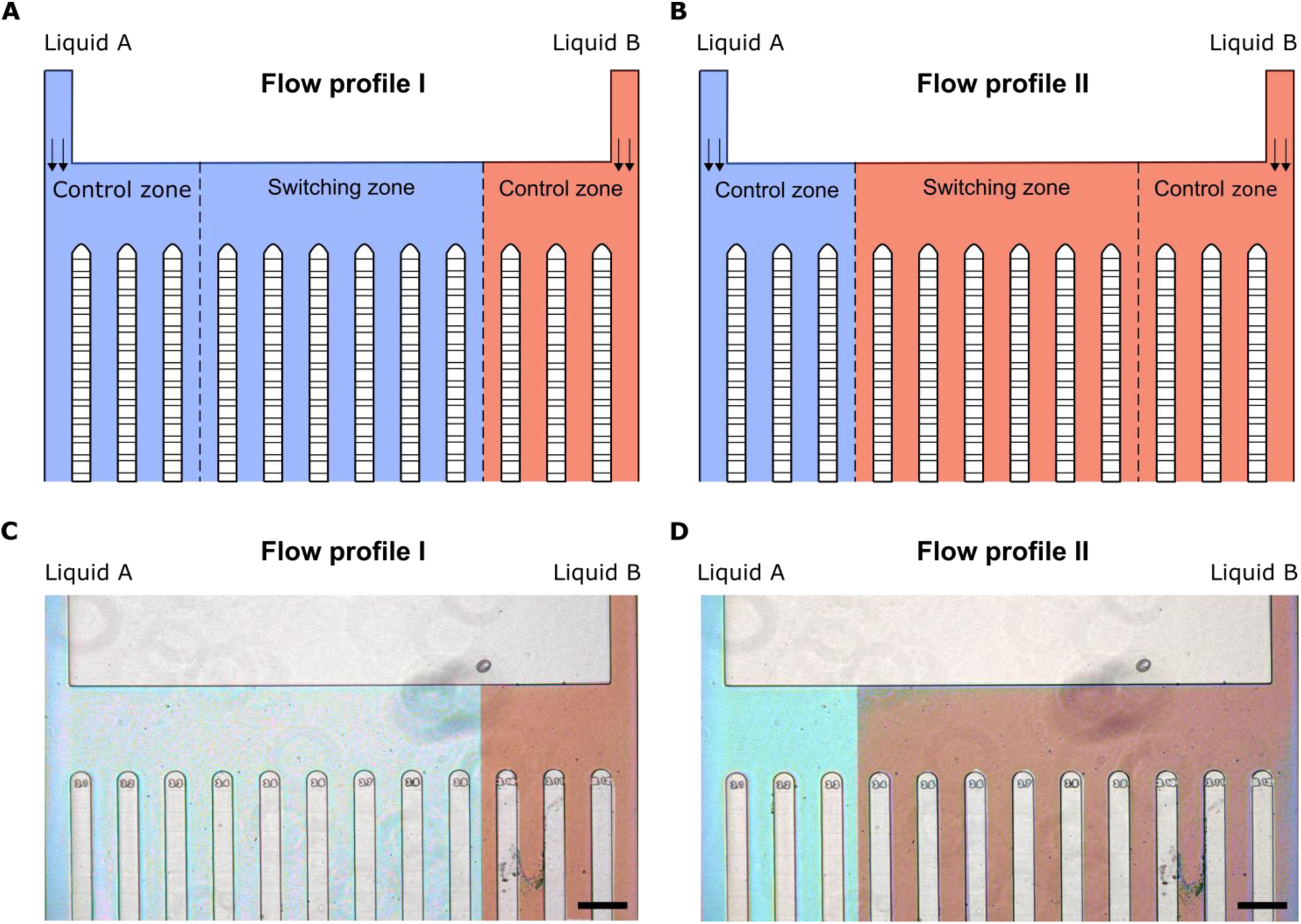
Illustration of the flow profile within the microfluidic dMSCC setup. A) Flow profile I: The six cultivation chamber arrays in the middle are flushed with liquid A. B) Flow profile II: The six arrays in the middle are flush withed the liquid B. C) and D) Microscopic images of the two flow profiles described in A and B. Scale bar 200 µm.

#### Validation of flow profiles and environmental conditions

Detailed flow experiments with fluorescein were performed to validate the setup and the environmental profiles that were obtained during dMSCC. The fluorescence signal of nine cultivation chambers in the different regions of the dMSCC setup (Fig. 3A) was analysed with an oscillation between fluorescein and ethanol in 1 minute intervals. The cultivation chambers C1-C3 were in the control zone I of the fluorescein, C4-C7 the switching zone and C8-C9 the control zone II, where no fluorescein was flushed. Fig. 3B shows three representatives cultivation chambers of each zone in the microfluidic device. During these 1-minute oscillation experiments, a fluorescence signal more than 95% is reached in the cultivation chamber after 5 seconds. Additional flow profile characterization experiments were performed with oscillation intervals of 5 seconds and confirmed these findings (see Fig. S2). Depending on the position of the cultivation chamber and thus the flow regime during the switching zone, the exchange rates of fluorescein can slightly differ. For example, cultivation chamber C4 (boundary between two zones) (see Fig. S3 – green curve) shows a faster exchange than the other selected cultivation chambers within the different regions in the arrays (see Fig. S3 – C1-C9). More detailed validation studies with oscillation times up to 0.5 seconds (2 Hz) were performed to test the limitation to realize stable and precise oscillating conditions. Fig. 3C sums up all validation measurements in a Bode plot (experimental results shown in black squares) with different oscillations times between 1 minute (0.017 Hz) and 0.5 second (2 Hz). Smaller oscillation intervals could not be performed due to the technical specification of the pumps. Here, the Bode plot^29^ was adapted from electrical engineering. The response is defined as normalized intensity of the fluorescence signal in the cultivation chamber. The plot shows, that medium switches up to 5 seconds can be performed, guaranteeing an almost complete medium exchange within the cultivation chamber. At oscillation intervals of 2 seconds (0.5 Hz) the fluorescence signal is 82% so that a full exchange of nutrients cannot be guaranteed anymore at this oscillation frequency. We conclude, for higher frequencies than 0.2 Hz, only a low medium exchange is feasible anymore. At very high oscillation intervals the system showed a transient effect (see Figure S4). At these oscillation frequencies, an average value was determined after the system had settled. At higher oscillation frequencies than 1 Hz, the signal in the cultivation chamber decreases only slightly until a stationary state of ∼ 50 % is reached.

**Fig. 3:**
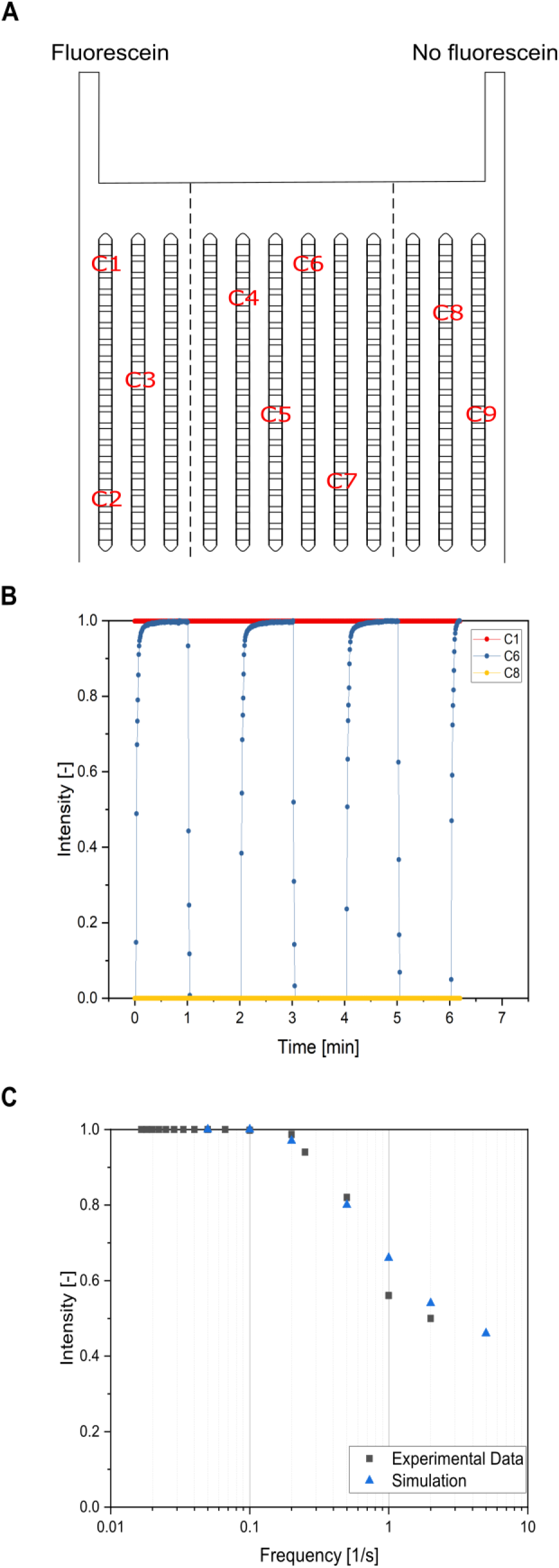
Experimental and model-based validation of substrate exchange within single-cell cultivation chambers. A) Schematic overview of the microfluidic device with selected cultivation chambers for the analysis of the fluorescence signal. B) Fluorescence profile of the 1 minute oscillation between fluorescein and ethanol. Here one cultivation chamber from each region in the microfluidic device is shown. Cultivation chamber C1 is in the control zone I (fluorescein region), C6 in the switching zone and C8 in the control zone II (no fluorescein region). C) Bode plot illustrating the fluorescence intensities in the cultivation chambers depending on different oscillation intervals of fluorescein from 100 s (0.01 Hz) to 0.2 s (5 Hz). The experimental validation (black squares) shows the fluorescence signal in cultivation chamber C6. The corresponding CFD simulated concentrations (blue triangles) are normalized by the maximum substrate concentration at the inlet.

CFD simulations were performed to verify the experimental findings (Fig. 3C – blue triangles). The measured substrate concentrations for each frequency are normalized, with the maximum substrate concentration measured in the cultivation chamber in steady state for the lowest frequency denoting 100%. The simulation verifies the experimental fluorescence results in the range 0.01 Hz to 2 Hz. Limitation of full medium exchange (∼ 97%) can be shown for intervals of 5 seconds (0.2 Hz), which is in good agreement with the respective experimental result. In the case of changes that are greater than 0.5 Hz, a slight offset of the simulation from the experiment can be observed. Above the limit of 0.5 Hz, the simulation shows a faster exchange of the medium compared to the experimental results. The following aspects can explain the discrepancy: First, the flow rate has an increasing impact on the exchange of substrate when oscillations get faster. The selected flow rate for the simulation that is based on an approximation may not perfectly reflect the flow rate in reality. Furthermore, the correlation between fluorescein signal and substrate concentration in the cultivation chamber is not impeccably linear, e.g. the height of the respective chamber region on the chip can influence the signal. However, the simulations confirm the operational limits of the dMSCC device.

The results show, that dMSCC up to 5 seconds (which corresponds well with environmental changes within bioreactors) can be performed with the presented setup, keeping in mind, that diffusivity of molecules can differ. Furthermore, cell growth within the cultivation chamber can influence diffusion time into the centre of the colonies/ chambers and can have an influence on the available concentration and thus on the substrate uptake.^30^

### Application – dMSCC of *Coryebacterium glutamicum*

#### Uniform oscillation profiles – Colony growth

The *C. glutamicum* strain WT ATCC 13032 is an important microbial host for industrial amino acid production^31–33^, such as L-lysine^34^ and L-glutamate^32^. In the first set of experiments, we investigated the microbial growth under oscillating (rectangular) feast and famine environmental conditions with different uniform oscillation times between 10 seconds and 1 hour (Fig. 4). Oscillation was carried out between the complex BHI medium and phosphate buffer (PBS) at 30 °C under aerobic conditions. The overall ratio between BHI medium (feast) and PBS (famine) was equal in all experiments.

**Fig. 4:**
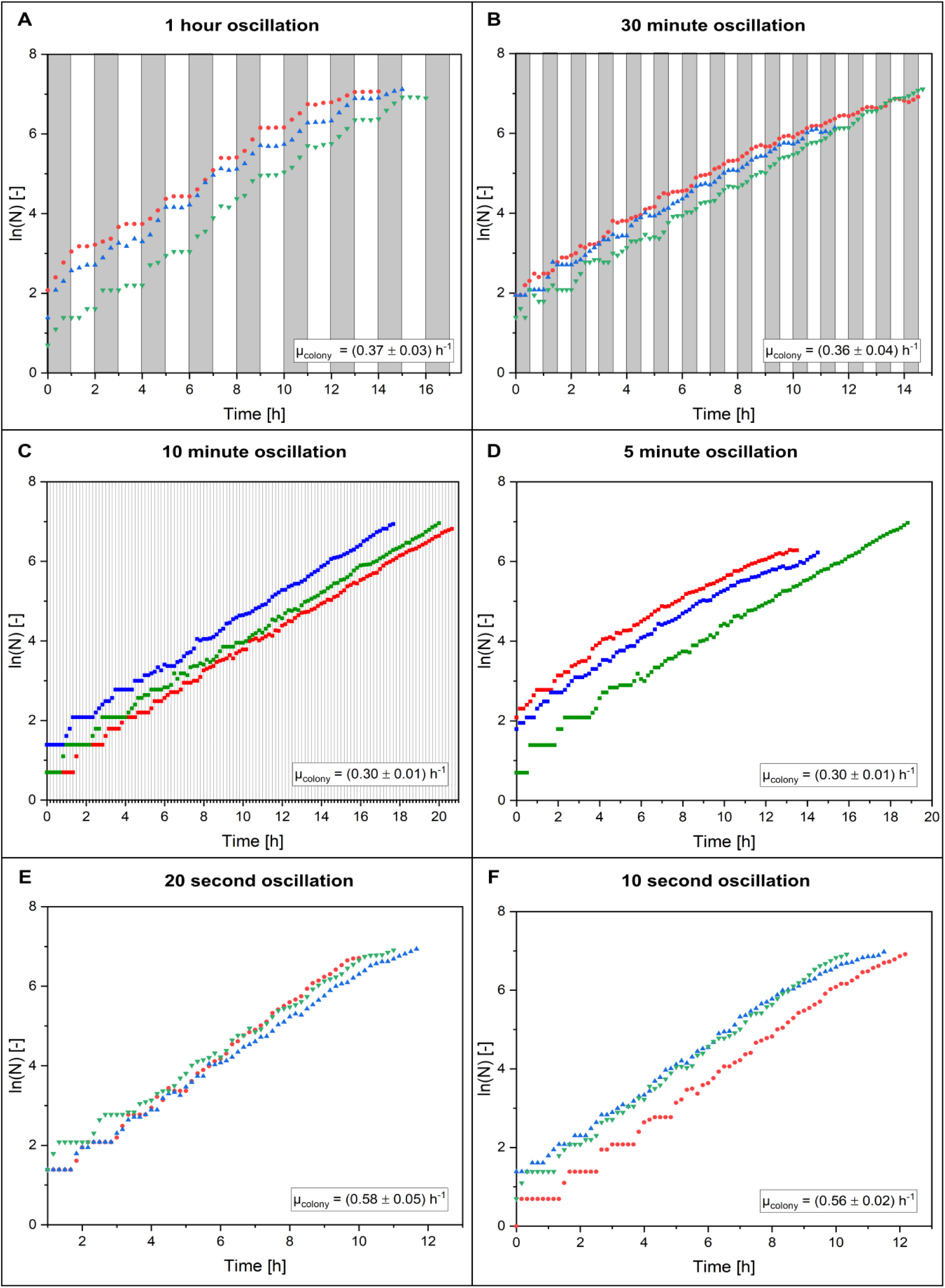
Growth curves at different oscillation frequencies between BHI medium and PBS buffer. The grey areas show the BHI medium pulses and the white areas the PBS buffer pulses. At higher oscillation frequencies the different pulse was not shown. A) 1 hour oscillation. B) 30 minute oscillation. C) 10 minute oscillation. D) 5 minute oscillation. E) 20 second oscillation. F) 10 second oscillation. Three colonies (red, green, blue) are shown.

As shown in Fig. 4A, biphasic growth behaviour can be observed at 1 hour oscillation cultivation. During BHI perfusion cells grow exponentially. During the intervals with PBS perfusion, colony growth was interrupted, and cells show no growth at all (Fig. 4A and see the ESI V1). The colony growth rate over the whole cultivation is µ_colony_ = (0.37 ± 0.03) h^−1^. Further experiments with higher oscillation intervals e.g. 45 minute show a similar growth behaviour (see Fig. S6 and the ESI V2). Here, a similar colony growth rate of µ_colony_ = (0.32 ± 0.03) h^−1^ was observed. A colony growth rate of µ_colony_ = (0.36 ± 0.04) h^−1^ was observed at the 30 minutes oscillations (Fig. 4B and see the ESI V3). At higher oscillation intervals of 5 min (0.003 Hz) to 15 min (0.001 Hz) a growth of µ_colony_ = (0.30 ± 0.01) h^−1^ was observed (Fig. 4C and D and see the ESI V4 – V6). Here, colonies show continuous growth in comparison to the low oscillating intervals, which is occasionally interrupted by a clear zero-growth (Fig. 4C and 4D and see Fig. S8 – S10). The 1 minute oscillations of BHI medium and PBS buffer show a faster growth as at the lower oscillation intervals (> 0.003 Hz). Here, a colony growth rate of µ_colony_ = (0.49 ± 0.07) h^−1^ was observed (see Fig. S10 and the ESI V7). For faster oscillation intervals of 20 s (0.05 Hz) a growth rate of µ_colony_ = (0.58 ± 0.05) h^−1^ was obtained (Fig. 4E and 4F and see the ESI V8 and V9). Under oscillating conditions, a positive influence of frequency on the growth rate was observed from a threshold of 1 minute (0.0167 Hz).

Several types of control experiments were performed with BHI and PBS buffer to guarantee technical functionality of the microfluidic device. The constant perfusion of the cells with BHI resulted in colony growth rates of µ_colony_ = (0.90 ± 0.04) h^−1^. To exclude that the pressure difference required for the oscillation of the medium has an influence on the colony growth a switch between BHI and BHI was performed in 10 second oscillation intervals, which resulted in a colony growth rate of µ_colony_ = (0.92 ± 0.02) h^−1^. These findings show that pressure variation which might result from the medium oscillations have no significant effect on the colony growth rate. As a third control experiment, BHI and PBS were mixed in a ratio of 1:1 to rule out any influence on growth and thus nutrient limitation which could happen by mixing the two components. The cells were continuously cultivated with the 1:1 BHI/PBS solution, achieving a colony growth of µ_colony_ = (0.90 ± 0.11) h^−1^, showing that a pure dilution of the BHI medium has also no effect on the growth rate and physiology of the cells during single-cell cultivation (see Fig. S14A-C).

All experiments performed under oscillating environmental conditions show a trend of a decrease in the colony growth rate compared to standard perfusion cultivations. An overview of the different colony growth rates is given in the frequency plot in Fig. 5. At low oscillation intervals (1h, 45min, 30 min), a colony growth rates of µ_colony_ = (0.32 – 0.37) h^−1^ are obtained. Cells respond to changes in the external environment by a complex regulatory system whose aim is the activation of transcription factors controlling the expression of a pool of genes necessary for the adaptation to the new environment.^35,36^ This leads to temporal starving environmental conditions and thus growth stop of all cells within the colony (µ_colony_ = (0.03 ± 0.02) h^−1^). All cells start to regrow after a medium switch to BHI (data not shown), indicating that cellular viability is maintained during the given starvation phases of each individual cell. The reduced growth rate of µ_colony_ ∼ 0.35 h^−1^ can be explained by a biphasic growth, with growth rates of µ_colony_ = (0.65 ± 0.03) h^−1^ during BHI pulse perfusion (non-continuous perfusion), and µ_colony_ ≈ 0 h^−1^ during PBS perfusion (see Fig. S15). We are currently not able to explain the differences in these growth rates µ_colony_ = (0.65 ± 0.03) h^−1^ vs. µ_colony_ = (0.90 ± 0.04) h^−1^ which are obtained during continuous BHI perfusion (see Fig. S14A). We assume, that temporal interruption of cell growth has a significant and temporal ongoing influence on the cellular metabolism and thus on the maximum obtainable growth rates during the BHI growth phases

**Fig. 5:**
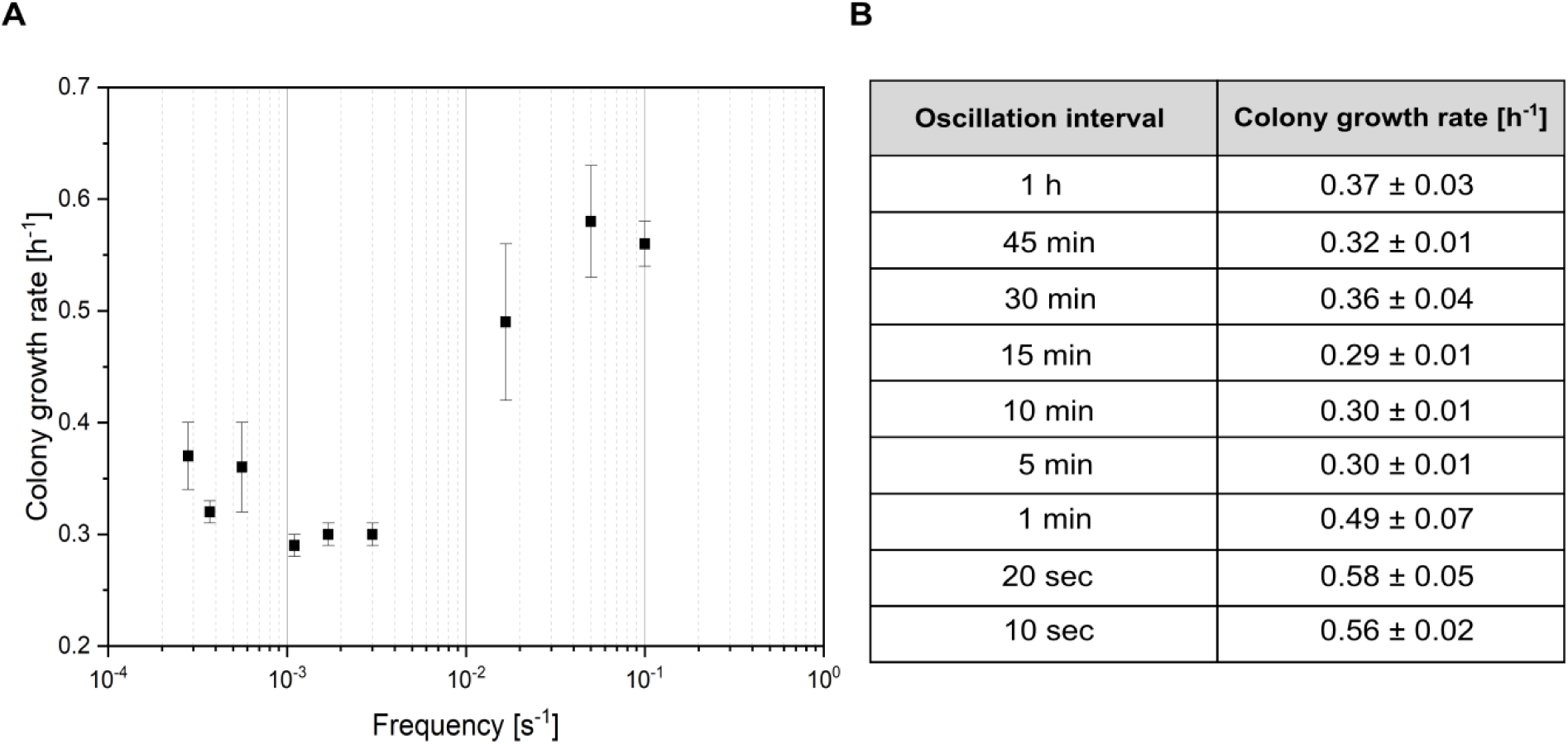
Overview of colony growth rates at varying oscillation frequencies. A) Frequency plot. B) Obtained colony growth rates at different oscillations intervals (N = 8 – 10).

At oscillation intervals between 5 and 15 minute growth rates show a further significant decrease of the growth rates of µ_colony_ = (0.29 – 0.30) h^−1^ with a (semi-) continuous growth profile. Some colonies show a continuous growth profile (see Fig. S9 – red), while in some colony’s distinct non-growth phases were observed (see Fig. S8 – red). Interestingly, the non-growth phases do not agree with the time-intervals of the medium switches. During 5 – 15 minute oscillations, the district non-growth phases show larger time-scale (∼ 45 min) (see Fig. S8). Currently, we have no detailed explanation for the observed growth phenomena. We suspect that the environmental oscillation in these frequencies have a significant effect onto the gene regulatory network of the cells and might thus result in irregular growth patterns. In future, detailed studies and analysis of single-cell behaviour might give further insights into this phenomenon. In comparison, during environmental oscillations at low frequencies, the regulatory network underlies no significant changes over the period of PBS perfusion. At high oscillation intervals (1 min, 20 sec, 10 sec) a continuous growth profile was measured, with a growth rate of µ_colony_ = (0.49 – 0.58) h^−1^. Cultivation with oscillations between 1 h and 5 min show a significant impairment of the colony growth rate compared to cultivations with high frequencies (1 min – 10 sec) and compared to BHI control experiment without switch (see Fig. S14A).

The reduced growth ranges at intermediate intervals (5 – 15 minute oscillation) cannot be explained conclusively at this time point. We assume, that oscillation intervals interfere with cellular regulation mechanisms which operate as a combination of different levels and stages of the flow of genetic information^37^, for example: mRNA degradation (∼ 10 min), gene splicing and translated protein (∼ one cell generation).^38,39^ These processes can only be carried out to a limited extent at high oscillation intervals since only limited new proteins can be encoded during the buffer pulses. There is a positive linear correlation between the RNA/ total protein ratio and the growth rate. More RNA is produced during faster growth, but the protein production rate remains constant. The oscillations can influence this ratio, which means that less RNA is produced and the ratio between RNA and total protein is smaller. The translation efficiency can be achieved by changing the nutrient quality of the medium, so that when the nutrient quality is low, translation is inhibited and the ratio of RNA to total protein is reduced.^40^

The growth rates of µ_colony_ = (0.49 – 0.58) h^−1^ found at high oscillation intervals (< 1 min) could be explained by differences in metabolism due to oscillations, leading to metabolic adaption rather than regulatory adaptation^41^. Adjustments in metabolism could lead to a change in the energy transformation ATP synthase.^42^ When cells use memory mechanisms as survival strategy for long-term cultivation, fitness costs resulting from gene regulatory expression may be reduced after the first stimulus is applied, as in the case of the carbon oscillation by Lambert et al.^22^. This effect can also occur here with the high oscillation frequencies, so that the cells do not have to constantly adapt to the new conditions, which increases the growth rate compared to the lower oscillation frequencies.

#### Uniform oscillation profiles – Single-cell length

Besides bulk parameters, single-cell parameters such as cell length and morphology can be analysed during dMSCC. Looking more detailed into the experimental results shown in the last section (colony growth behaviour), single-cell length distributions were analysed. Fig. 6 shows the cell size distributions for six oscillation frequencies for three colonies with approximately 150 cells after 12 hours of cultivation. There is a large variation of the average and width of the population depending on the oscillation frequency. At the 1 hour oscillation the average cell size of (3.4 ± 0.1) µm was observed. At shorter starvation periods the average cell size decreased so that the 45 minutes oscillation an average cell size of (2.2 ± 0.1) µm, the 30 minutes oscillation an average cell size of (2.0 ± 0.1) µm, the 10 minutes oscillation an average cell size of (2.6 ± 0.1) µm and the 10 seconds oscillation an average cell size of (2.5 ± 0.2) µm were observed (see Fig. S16). The cell length measurement was independent of the choice of the cultivation time, before or after the pulse (see Fig. S17).

**Fig. 6:**
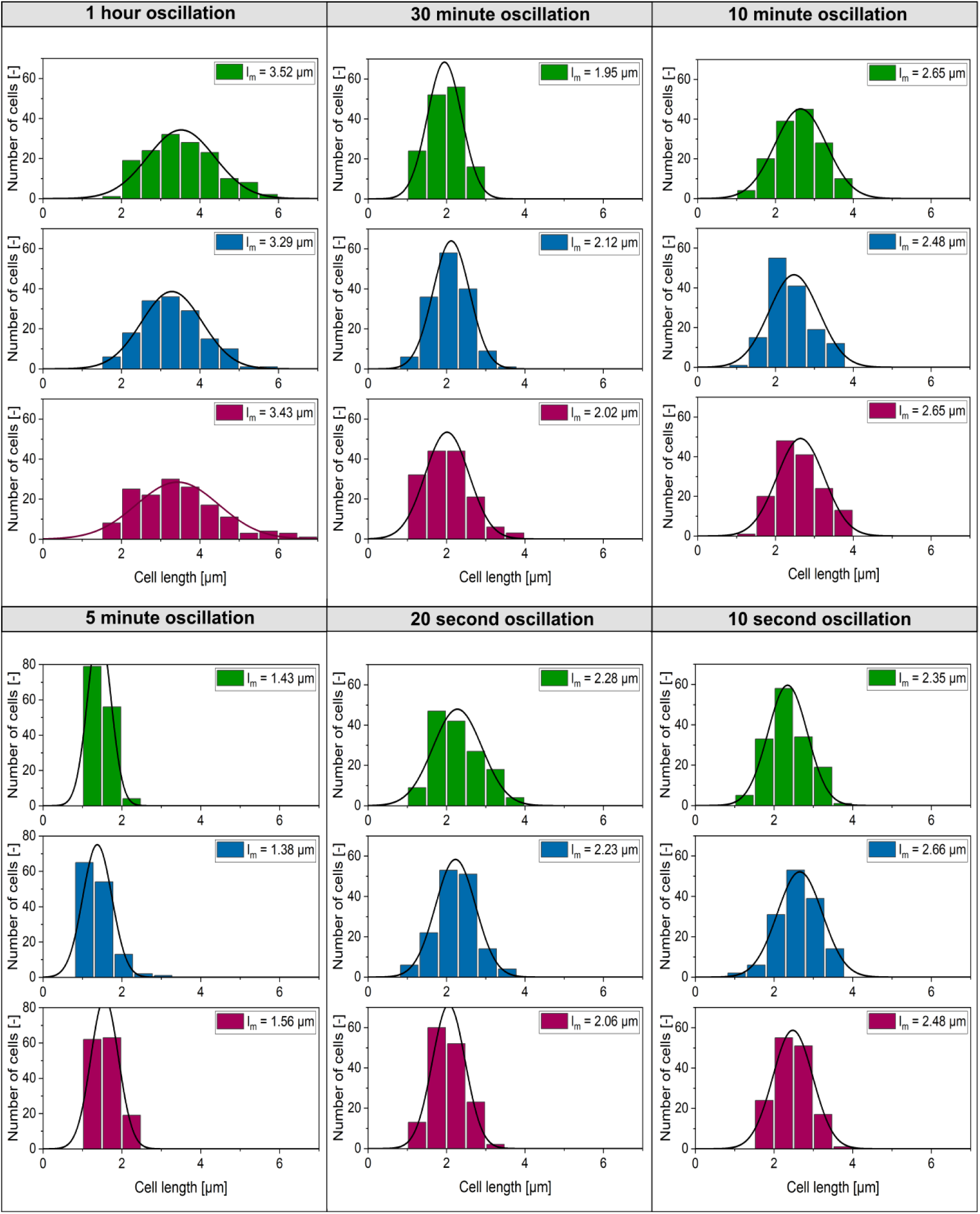
Cell length histogram of the dMSCC with different oscillations frequencies between BHI medium and PBS after 12 h of cultivation, three colonies with N = 150 (green, blue, red).

There is a positive trend between colony growth rate and average cell length. High growth rates lead to significant increased single-cell cell lengths (Fig. 7). Only the cell length of the one-hour oscillation is outside the linear trend. This correlation between the growth rate and cell size was firstly observed for *Salmonella typhimurium* organisms under batch conditions by Schaechter et al.^43^.

**Fig. 7:**
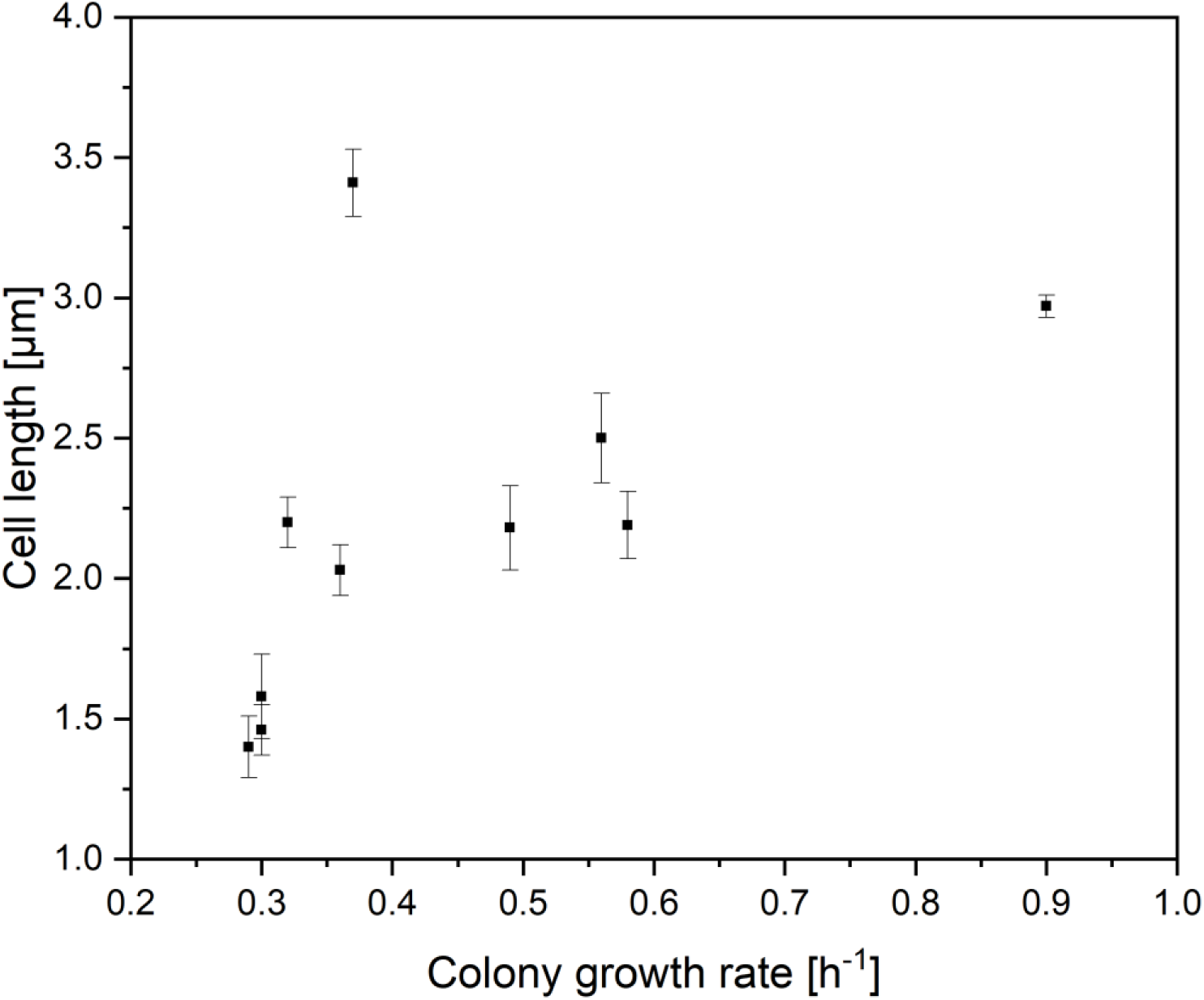
Colony growth rate from the oscillation cultivation (seen in Fig. 4) as a function of the cell length.

#### Irregularly oscillating profiles

The dMSCC setup was designed so that medium can be oscillated at high frequencies and with varying lengths (amplitude is constant). To test this feature, first experiments for the reproduction of simulated cellular lifelines^4^ of nutrient gradients in large scale bioreactors were performed. In comparison to bioreactor gradients, which are complex and arbitrary and not completely understood in the effects of a single cell, here irregular random transitions between feast (BHI-flushed) and famine (PBS-flushed) conditions were performed. We created irregular oscillations with times between 5 seconds and 1 minutes in an coarse-grained analogy of the simulation of lifelines in large scale bioreactors from Haringa et al.^4^ (see Fig. S18). The irregular oscillations were varied in the medium supply, the length of PBS buffer pulses was kept constant in all experiments (Fig. 8). Three experiments were performed with different overall medium supply, the first experiment with 40% overall medium supply, the second experiment with 60% overall medium supply and the third experiment with 80% overall medium supply during the whole cultivation. The experiments show that the total amount of BHI has a positive influence on the growth rate. We found an exponential correlation (R=0,9999) between the colony growth rate and the total medium supply (see Fig. S19). For 80% overall medium supply over the whole experiment, an overall growth rate of µ_colony_ = (0.66 ± 0.03) h^−1^ is shown in Fig. 8C. In experiments with less overall medium supply, an overall growth rate of µ_colony_ = (0.51 ± 0.06) h^−1^ at 60% overall medium supply (Fig. 8B) and an overall growth rate of µ_colony_ = (0.42 ± 0.04) h^−1^ at 40% overall medium supply were obtained (Fig. 8A). The results are in agreement, with the results obtained in the uniform oscillation experiments with sub-minute resolution (µ ∼ (0.49 – 0.58) h^−1^, see Fig. 5), which are performed with a medium supply of 50% during the cultivation time.

**Fig. 8:**
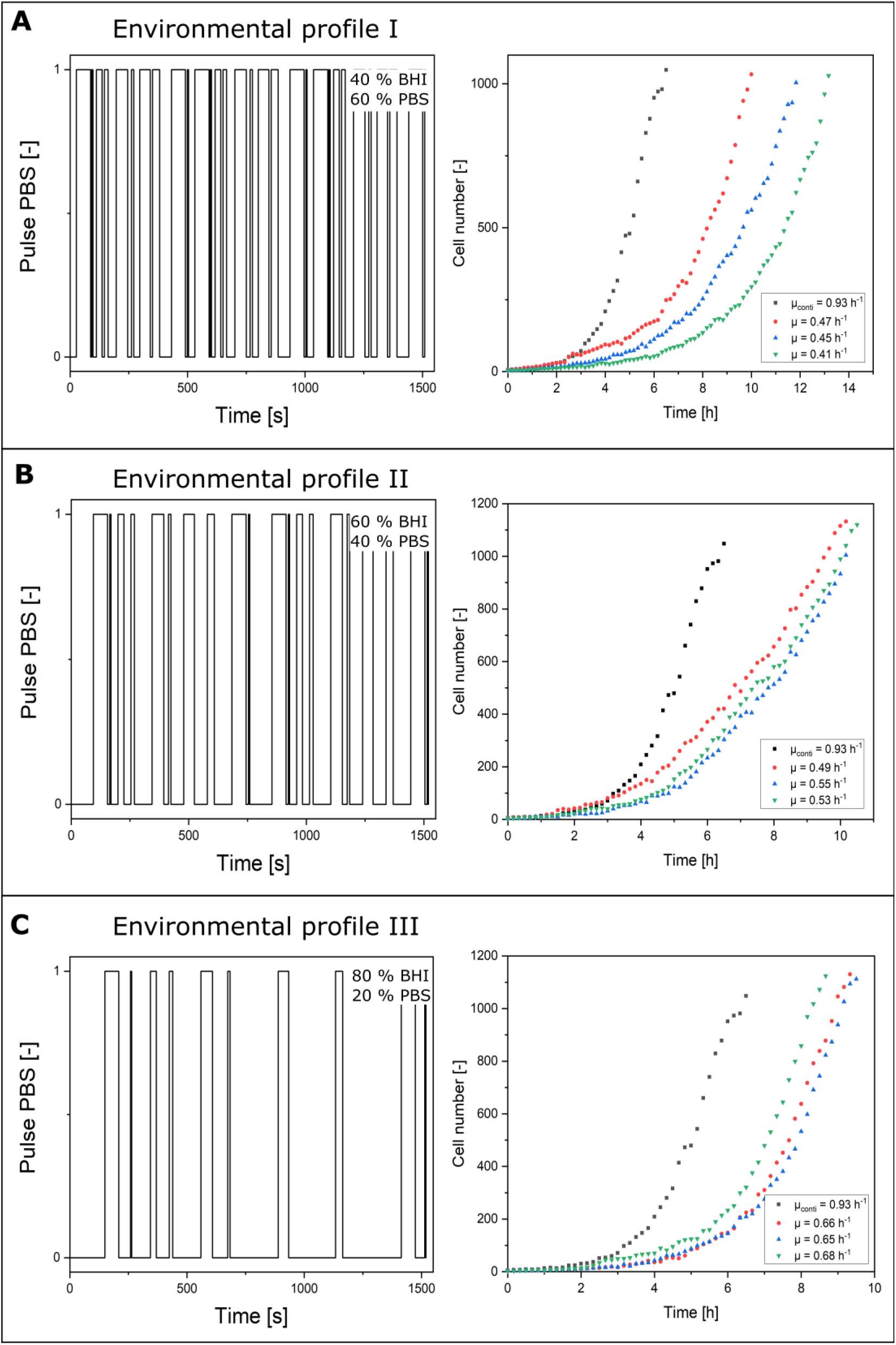
Feed profiles and measured overall colony growth rates for approximated lifelines with A) 40%, B) 60%, and C) 80% overall medium supply.

This lays the foundation for detailed studies of lifelines in bioreactors and the analysis of cellular behaviour under complex fluctuating environmental conditions, e.g., pH value, carbon source and general medium composition.^4^ The profiles need to be further optimized, since so far only on/ off profiles can be investigated. Different concentrations/ amplitudes can be simulated with a fast switching back and forth in the sub-second range, when the bacterial cells can perceive the difference between constant conditions and fast changes. In this case any concentration can be simulated by very fast oscillation. Furthermore, microbial interactions can be investigated under oscillating nutrient supply and different pH values.^44^

## Conclusions and Outlook

In this work, a new microfluidic single-cell cultivation setup for the cultivation and analysis of microbial single cells at dynamic environmental conditions (dMSCC) with high spatio-temporal resolution was developed. This setup enables fast exchange of medium conditions (in seconds range) as well as simultaneous control measurements of the medium conditions on a chip. Systematic experiments with *C. glutamicum* as a model system were performed with different fast (BHI) and famine (PBS) oscillation intervals between 1 hour and 10 seconds. Our experiments show that cellular growth behaviour was significantly affected at the different oscillating conditions. At low oscillation frequencies (2.7 · 10^−4^ Hz) a colony growth rate of µ_colony_ ∼ 0.3 h^−1^ was achieved. As the oscillation frequency increased (0.017 Hz), an increase in growth rates was observed (µ_colony_ ∼ 0.58 h^−1^). Additionally, we demonstrated for the first time the cultivation of microbial cells under randomly generated nutrient profiles similar to the environmental conditions present in large-scale bioreactors. With this new system in hand it will be possible to investigate the influence of different fast famine cycles, different media composition and concentrations on the cellular behaviour at the level of individual microbial cells. This lays the foundation of systematic studies of microbial behaviour at various oscillating environments, both in fundamental and applied microbiology and biotechnology.

## Author contributions

Conceptualization and investigation: ST, CG and AG. Formal analysis and validation: ST, PH, CG and AG. Visualization: ST and PH. Resources: EL and AG. Supervision: AG. Writing: ST, AG and PH.

## Conflict of interest

There are no conflict of interest to declare.

## Acknowledgement

Parts of this work were performed at the cleanroom facilities of the Department of Biophysics and Nanoscience as well as the Department for Physics of Supramolecular Systems and Surfaces at Bielefeld University. The authors would like to thank for all the help and support.

## Notes

### Competing Interest Statement

The authors have declared no competing interest.

