## Supplement Videos for "dMSCC: A microfluidic platform for microbial single-cell cultivation under dynamic environmental medium conditions"

**V1**

Growth of *C. glutamicum* in dMSCC at 1 hour oscillation between BHI medium and PBS buffer.

**V2**

Growth of *C. glutamicum* in dMSCC at 45 minute oscillation between BHI medium and PBS buffer.

**V3**

Growth of *C. glutamicum* in dMSCC at 30 minute oscillation between BHI medium and PBS buffer.

**V4**

Growth of *C. glutamicum* in dMSCC at 15 minute oscillation between BHI medium and PBS buffer.

**V5**

Growth of *C. glutamicum* in dMSCC at 10 minute oscillation between BHI medium and PBS buffer.

**V6**

Growth of *C. glutamicum* in dMSCC at 5 minute oscillation between BHI medium and PBS buffer.

**V7**

Growth of *C. glutamicum* in dMSCC at 1 minute oscillation between BHI medium and PBS buffer.

**V8**

Growth of *C. glutamicum* in dMSCC at 20 second oscillation between BHI medium and PBS buffer.

**V9**

Growth of *C. glutamicum* in dMSCC at 10 second oscillation between BHI medium and PBS buffer.
