## Supplement information for "dMSCC: A microfluidic platform for microbial single-cell cultivation under dynamic environmental medium conditions"

### SI Materials and methods

#### *CFD*

#### *Geometry and operating conditions*

The geometry was adapted from Probst et al.[1]. It has a symmetric design with two supply channels flanking the cultivation chamber. The cultivation chamber was modelled with the exact dimensions shown here, 90 µm width, 80 µm length and 0.8 µm height. It is directly connected to the supply channels that are 100 µm wide, 140 µm long and 10 µm high. Both supply channels are modelled 30 µm upstream and downstream of the cultivation chamber. Inlet and outlet boundary conditions were applied at these positions. Symmetry along a symmetry plane through the cultivation chamber was exploited to reduce computation time.

#### *Simulation setup*

Calculations of the stationary velocity field and the transient mass transport were performed using COMSOL Multiphysics 5.4.0.225 (COMSOL AB, Sweden). For the stationary velocity field, the stationary Navier-Stokes equations[2] for an incompressible, isothermal, Newtonian fluid were solved:

$$\mu \nabla^2 \boldsymbol{u} = \nabla p$$

$$\nabla \cdot \boldsymbol{u} = 0$$

where $\boldsymbol{u}$ denotes the fluid velocity vector in m $s^{-1}$, p the fluid pressure in Pa, and $\mu$ the dynamic viscosity in Pa s. The properties of the fluid within the channels and the cultivation chamber were treated as identical to pure water. The parabolic velocity profile at the inlets of the supply channels was applied using the laminar inflow feature of COMSOL Multiphysics with an inflow length of 100 µm. The flow rate at the inlet was set to be 200 nl $min^{-1}$. For the PDMS and glass walls, the no-slip condition was used.

For the mass transport the diffusion-advection equation according to Deen[2] was applied:

$$\frac{\partial c}{\partial t} + \nabla \cdot (-D\nabla c) + \boldsymbol{u} \cdot \nabla c = 0$$

where $c$ denotes the substrate concentration in mmol $L^{-1}$, $D$ the binary diffusion coefficient of the solute in water in $m^2$ $s^{-1}$, and $\boldsymbol{u}$ the velocity vector in m $s^{-1}$. The diffusion coefficient of glucose in water is $5.4 \cdot 10^{-10}$ $m^2$ $s^{-1}$.[3] A rectangular pulse function with maximum concentration of 222 mmol $L^{-1}$ and a fixed frequency between 0.05 Hz and 5 Hz was imposed as inlet concentration.

A frequency response analysis was conducted by comparing the normalized substrate concentrations measured in the cultivation chamber at a stationary state, with the maximum substrate concertation measured in total denoting 100%. A stationary state was assumed when the amplitudes measured in the cultivation chamber varied less than 1% over five consecutive oscillations. The substrate concentrations in the cultivation chamber were determined using the domain probe feature of COMSOL Multiphysics. The results are presented in a Bode plot.

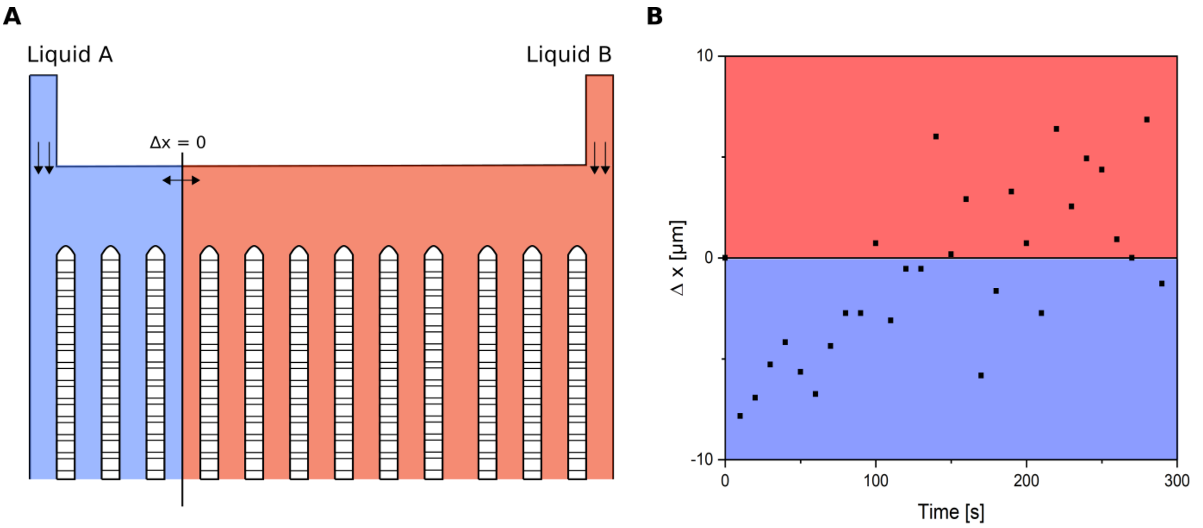

**Fig. S1**: Displacement of the boundary line between two laminar flows from set point ($\Delta x = 0$) at the 10 second oscillation.

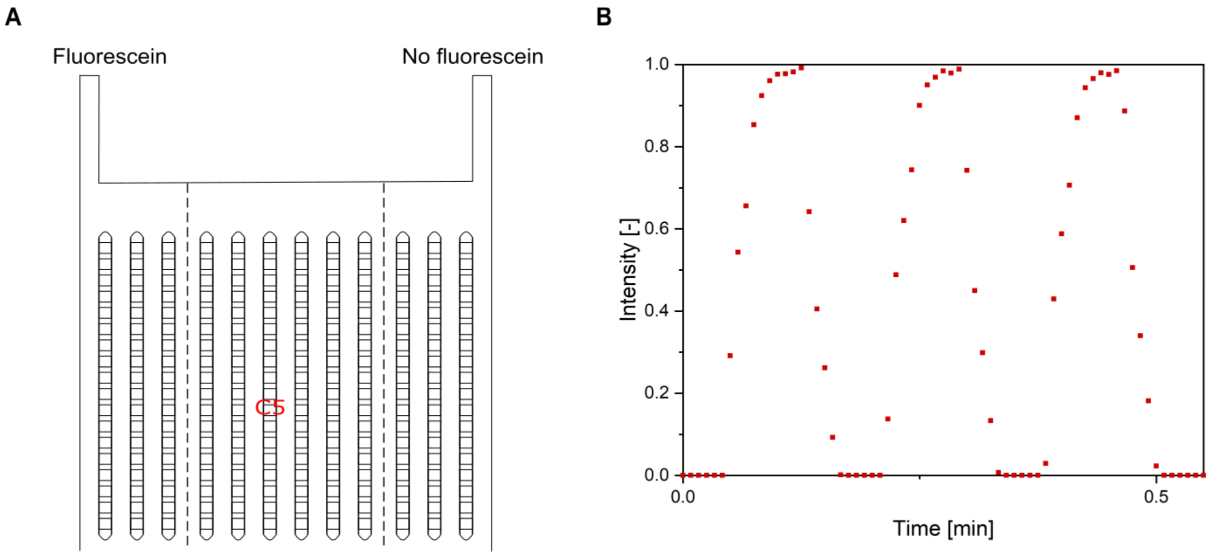


**Fig. S2:** Result of the experimental validation of the microfluidic device of the 5 second oscillation with fluorescein and ethanol at the position C6.

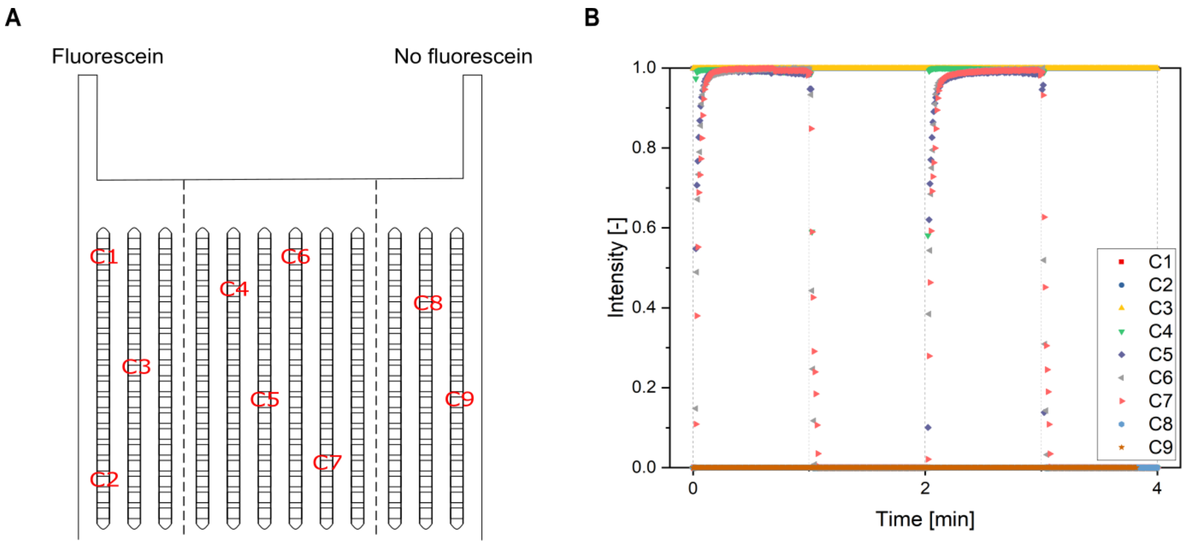


**Fig. S3**: Experimental validation of the microfluidic device. A) Schematic overview of the microfluidic device with selected cultivation chambers for the analysis of the fluorescence signal. B) Result of the 1 minute oscillation with fluorescein and ethanol for different positions.

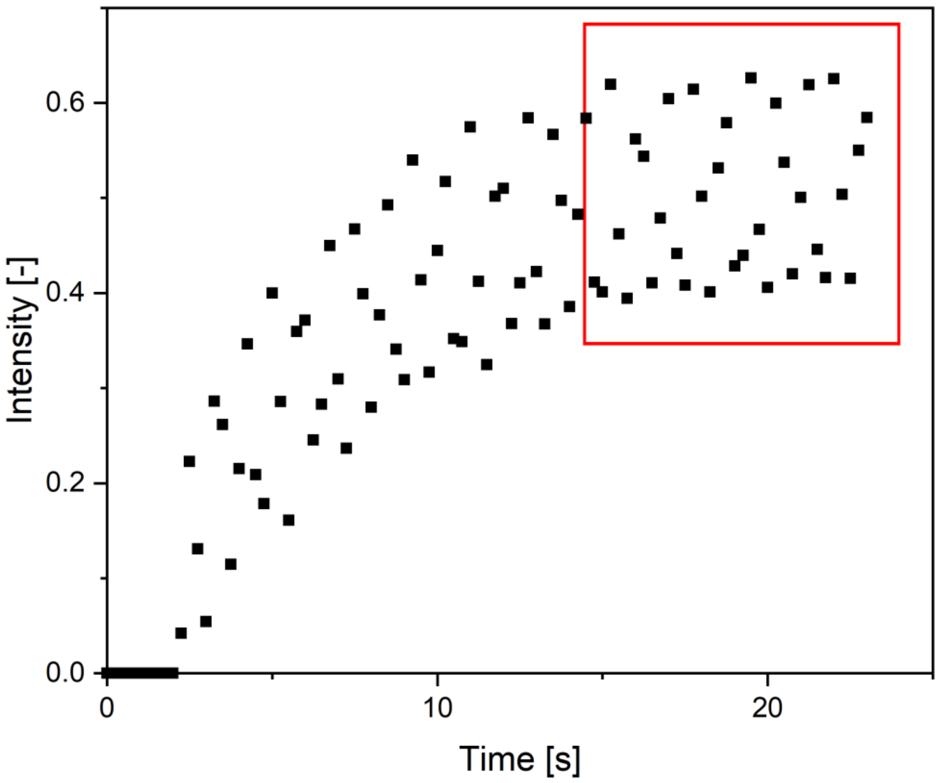


**Fig. S4**: Transient effect during the 500 ms oscillation. In the red area the average value after the transient oscillation was calculated

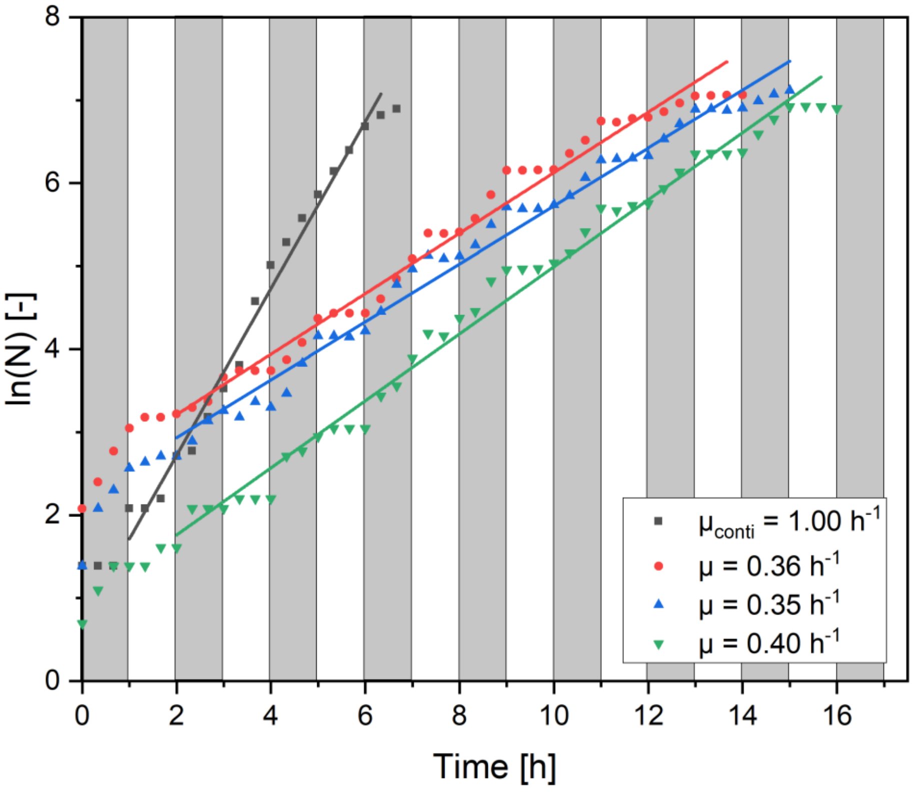


**Fig. S5**: Growth curves of dMSCC at 60 minute oscillation between BHI medium and PBS buffer with linear regression for the determination of the growth rate ($\mu_{colony}$ = (0.37 ± 0.03) $h^{-1}$). The grey areas show the BHI medium pulses and the white areas the PBS buffer pulses.

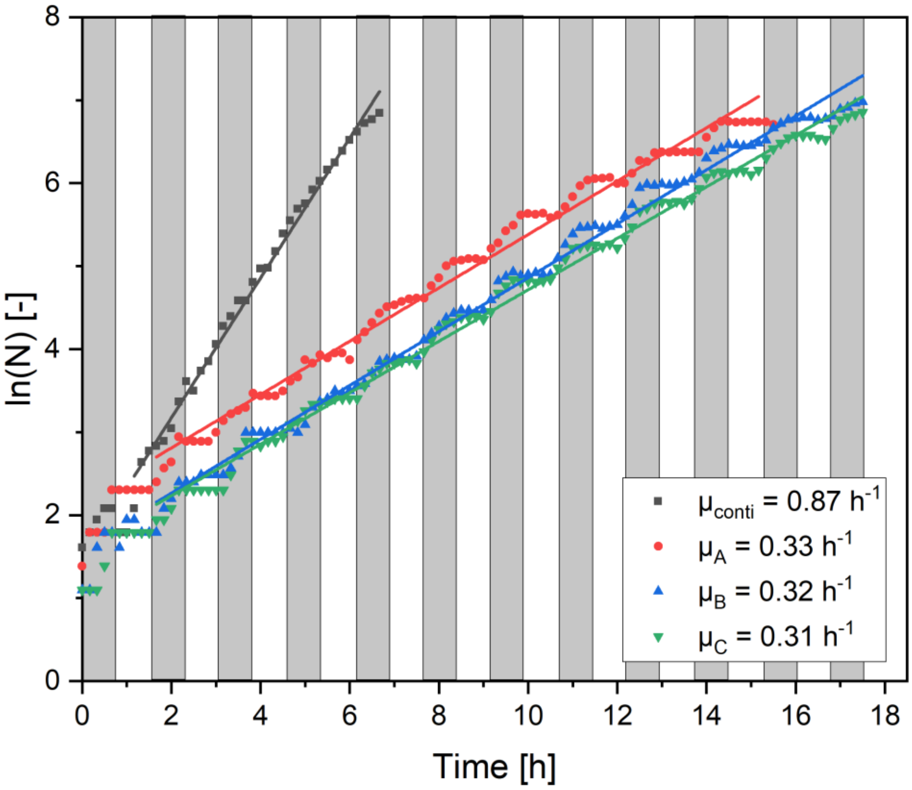


**Fig. S6**: Growth curves of dMSCC at 45 minute oscillation between BHI medium and PBS buffer with linear regression for the determination of the growth rate ($\mu_{colony}$ = (0.32 ± 0.01) $h^{-1}$). The grey areas show the BHI medium pulses and the white areas the PBS buffer pulses.

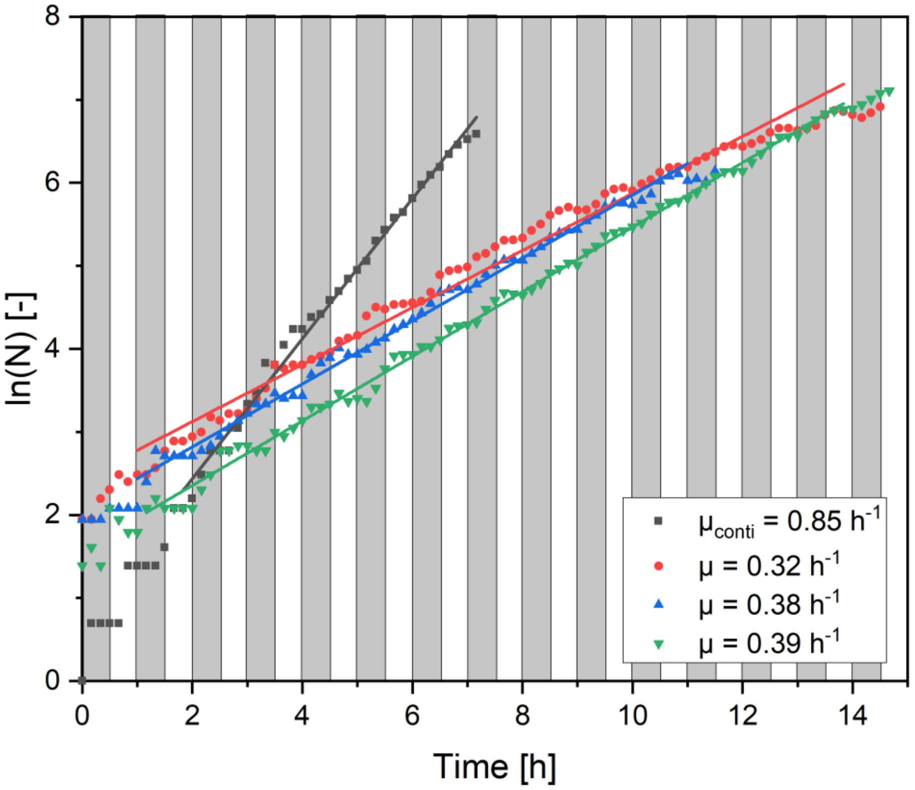


**Fig. S7**: Growth curves of dMSCC at 30 minute oscillation between BHI medium and PBS buffer with linear regression for the determination of the growth rate ($\mu_{colony}$ = (0.36 ± 0.04) $h^{-1}$). The grey areas show the BHI medium pulses and the white areas the PBS buffer pulses.

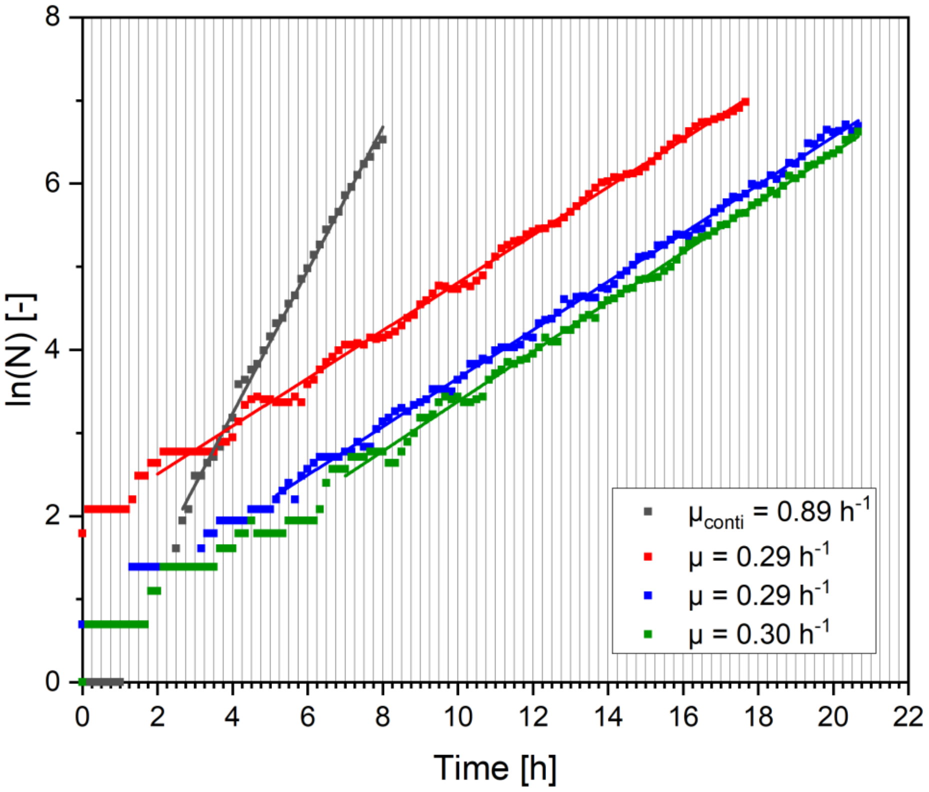


**Fig. S8**: Growth curves of dMSCC at 15 minute oscillation between BHI medium and PBS buffer with linear regression for the determination of the growth rate ($\mu_{colony}$ = (0.29 ± 0.01) $h^{-1}$).

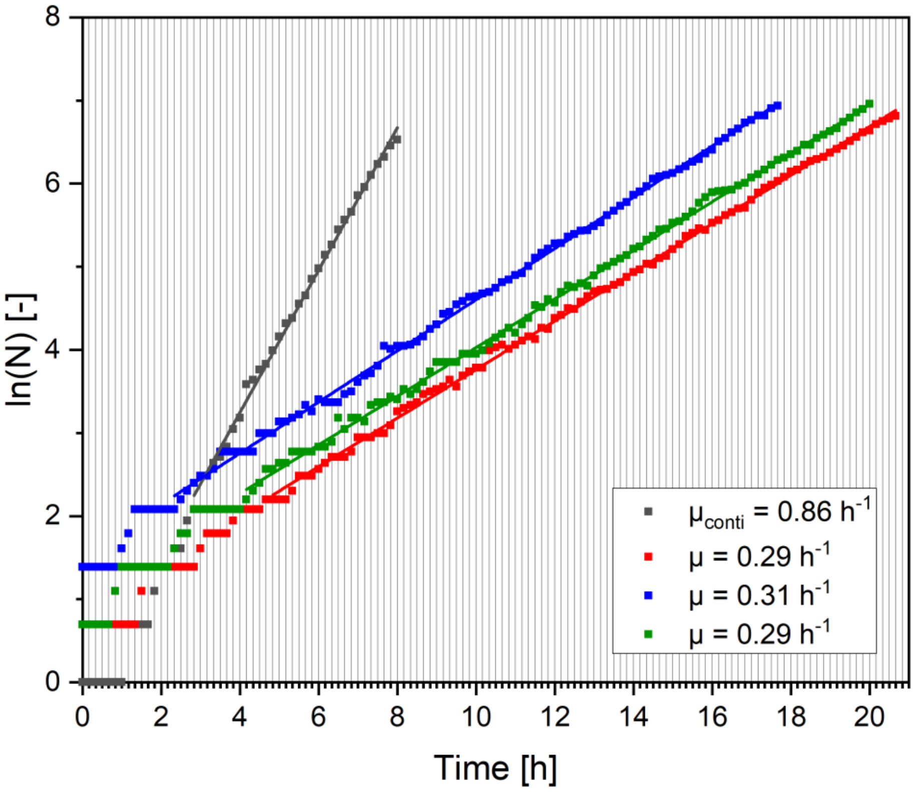


**Fig. S9**: Growth curves of dMSCC at 10 minute oscillation between BHI medium and PBS buffer with linear regression for the determination of the growth rate ($\mu_{colony}$ = (0.30 ± 0.01) h$^{-1}$).

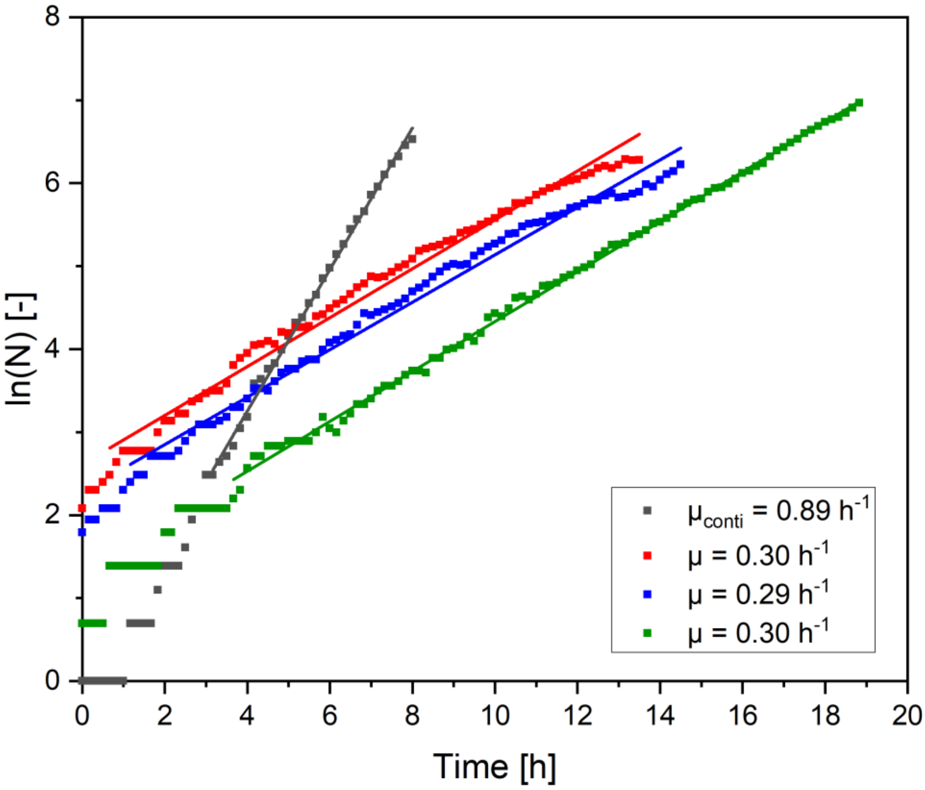


**Fig. S10**: Growth curves of dMSCC at 5 minute oscillation between BHI medium and PBS buffer with linear regression for the determination of the growth rate ($\mu_{colony} = (0.30 \pm 0.01)\ h^{-1}$).

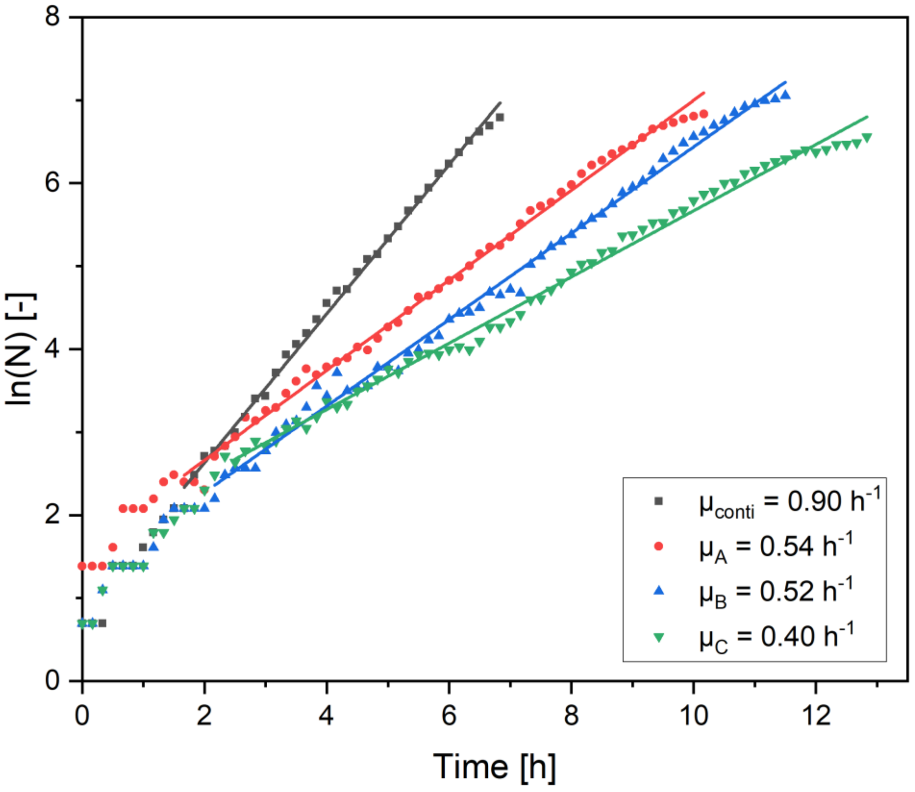


**Fig. S11**: Growth curves of dMSCC at 1 minute oscillation between BHI medium and PBS buffer with linear regression for the determination of the growth rate ($\mu_{colony}$ = (0.49 ± 0.07) $h^{-1}$).

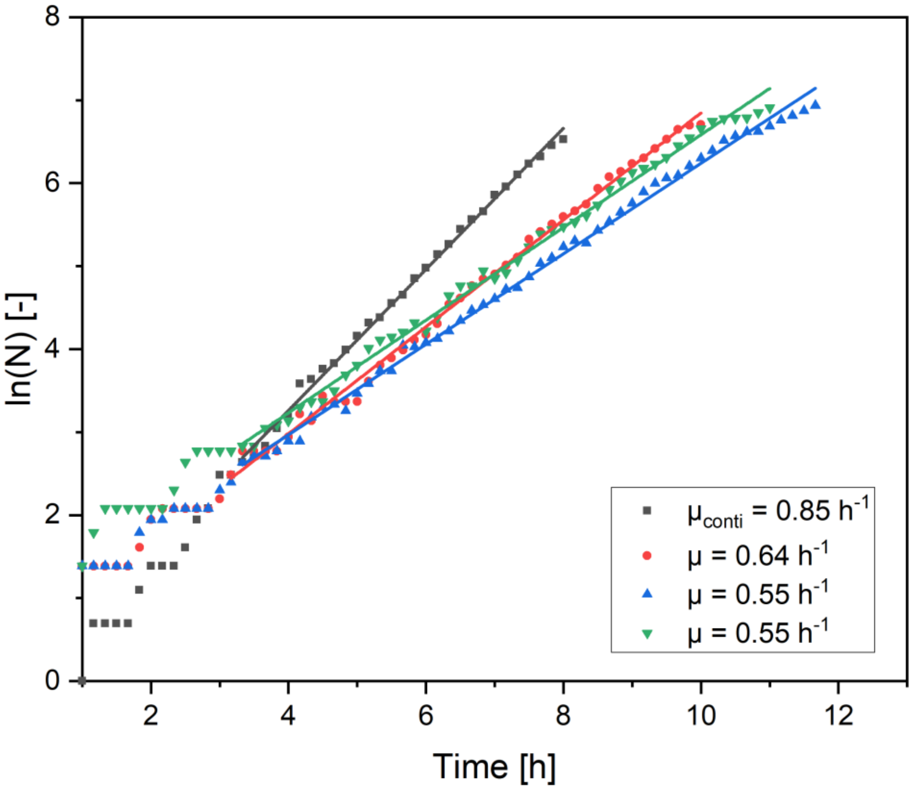


**Fig. S12**: Growth curves of dMSCC at 20 second oscillation between BHI medium and PBS buffer with linear regression for the determination of the growth rate ($\mu_{colony} = (0.58 \pm 0.05)\ h^{-1}$).

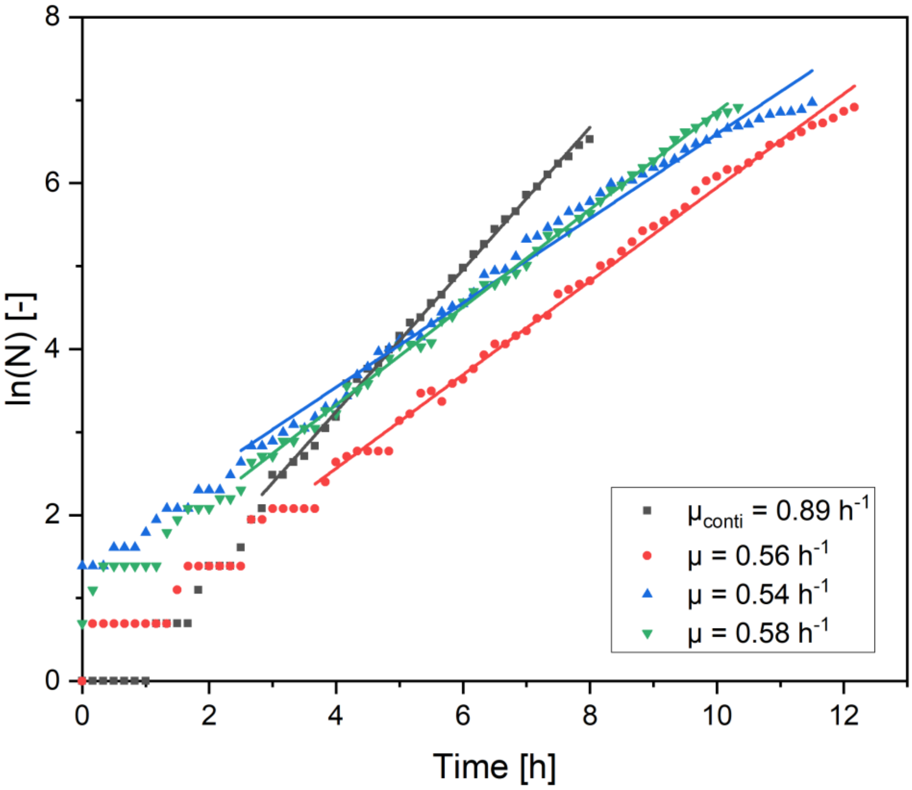


**Fig. S13**: Growth curves of dMSCC at 10 second oscillation between BHI medium and PBS buffer with linear regression for the determination of the growth rate ($\mu_{colony}$ = (0.56 ± 0.02) $h^{-1}$).

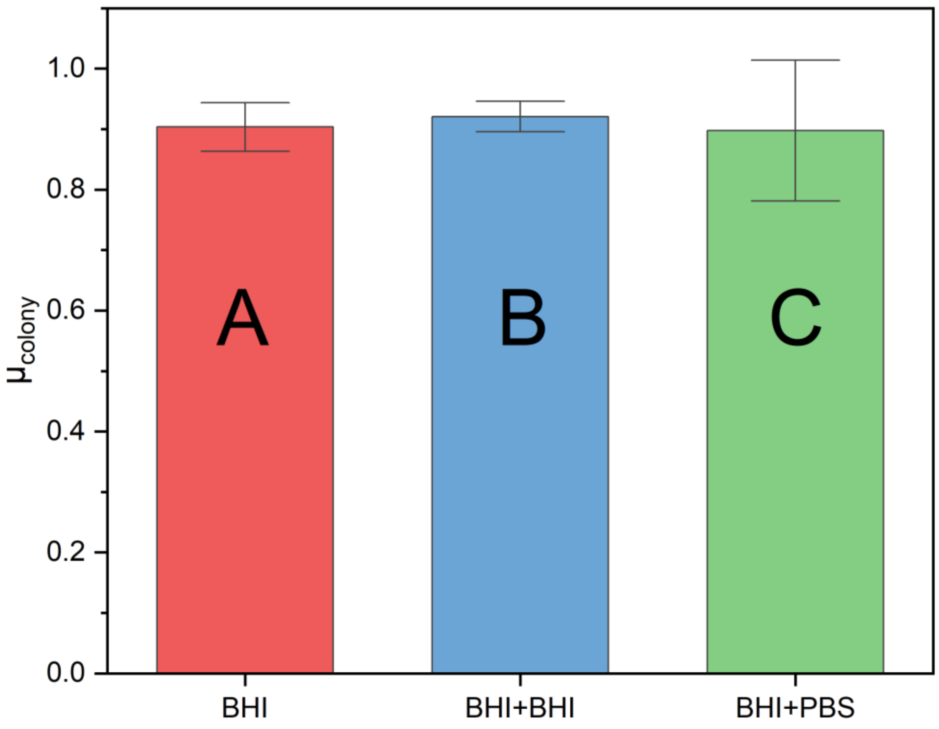


**Fig. S14**: Overview of different control experiments. A) Continuous cultivation in BHI medium (red). B) 10 second oscillation between BHI medium and BHI medium (blue). C) Continuous cultivation with a 1:1 mixture of BHI medium and PBS buffer (green).

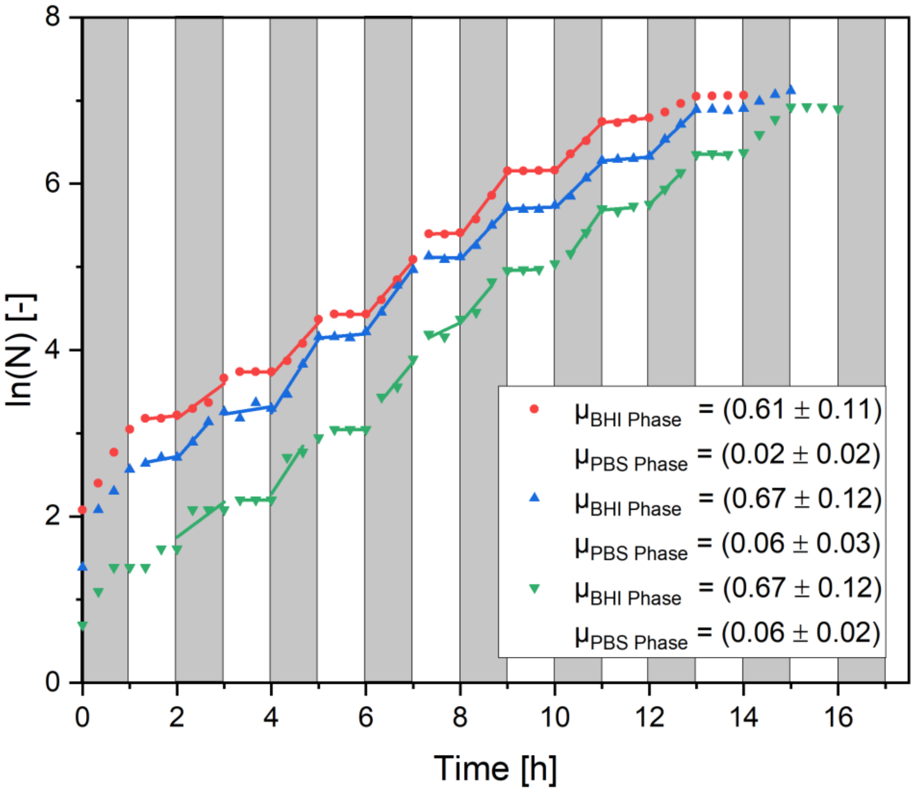


**Fig. S15**: Growth curves of dMSCC at 1 hour oscillation between BHI medium and PBS buffer with linear regression for the determination of the growth rate during the BHI perfusion phase and PBS perfusion phase.

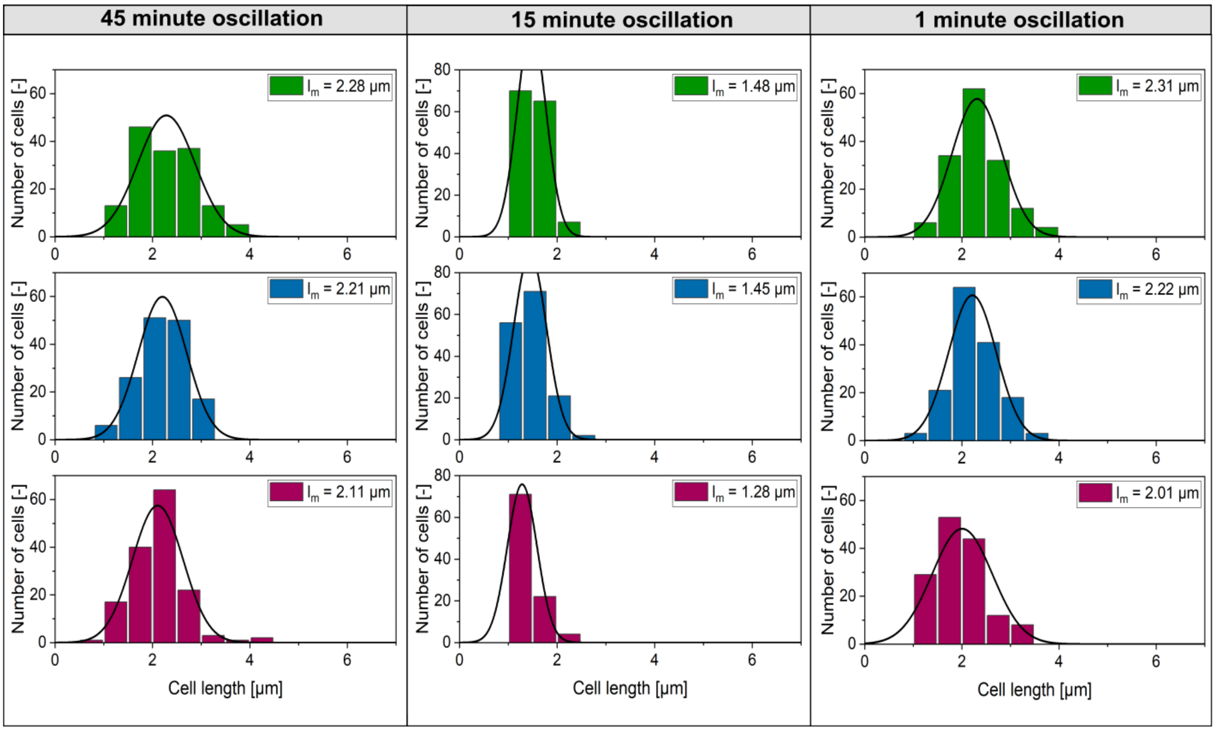


**Fig. S16**: Cell length histogram of the dMSCC with different oscillations frequencies between BHI medium and PBS buffer after 12 h of cultivation, three colony with N = 150 (green, blue, red).

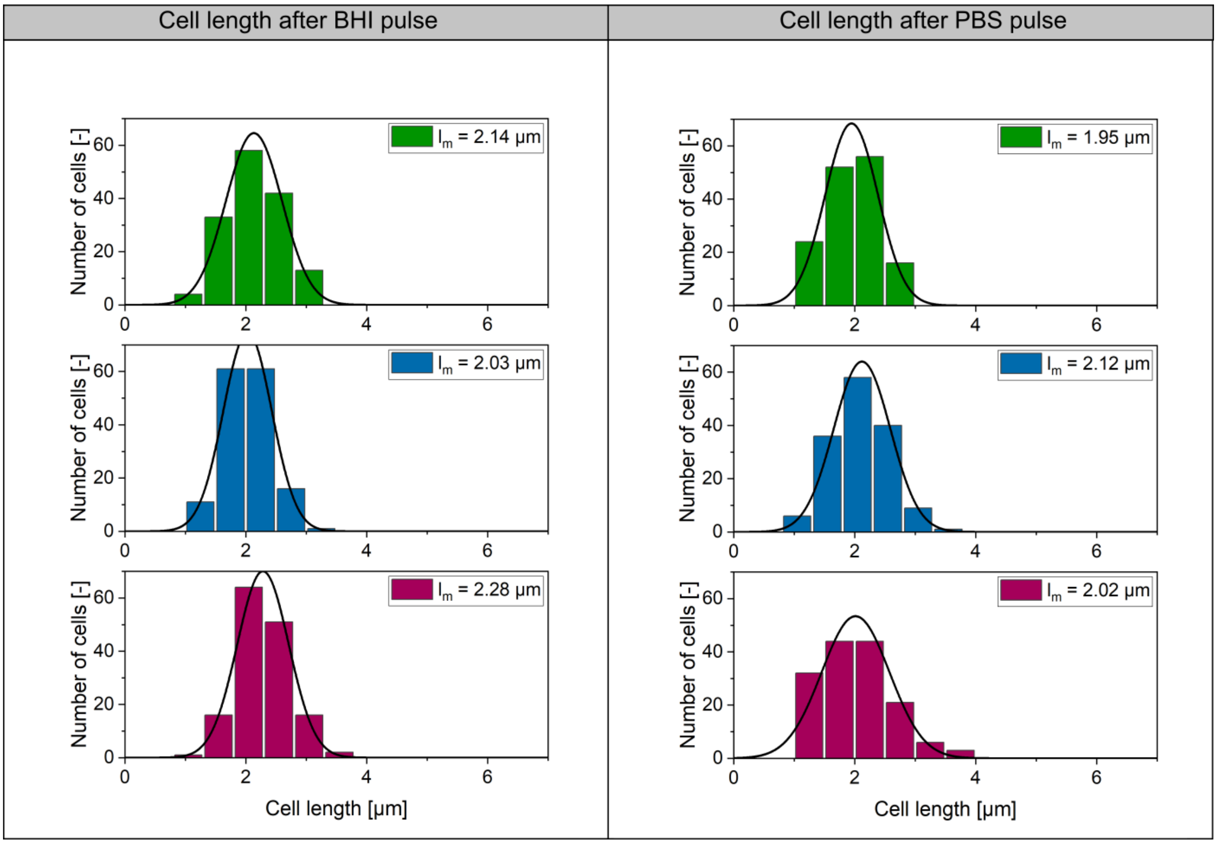

**Fig. S17**: Cell length histogram of the dMSCC for the 30 minute oscillation after BHI pulse and after the PBS pulse after 12 h of cultivation, three colony with N = 150 (green, blue, red).

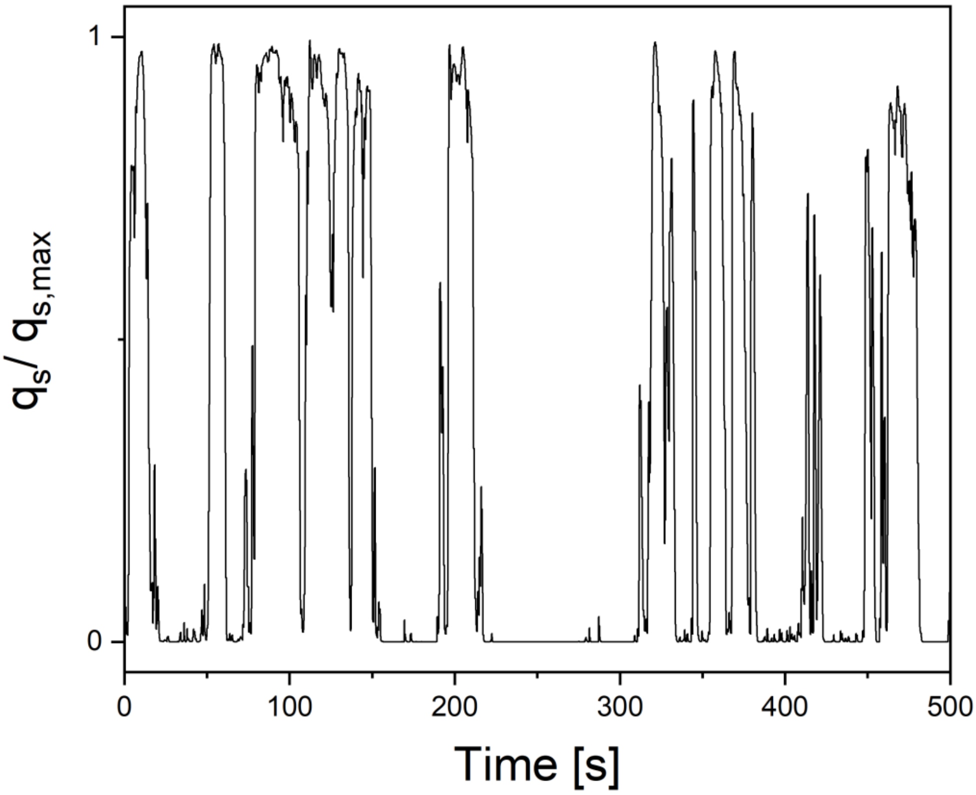


**Fig. S18**: Typical lifeline of a large-scale bioreactor. Copyright 2020, from Biochemical Engineering Journal. Adapted with permission from Haringa et al.[4].

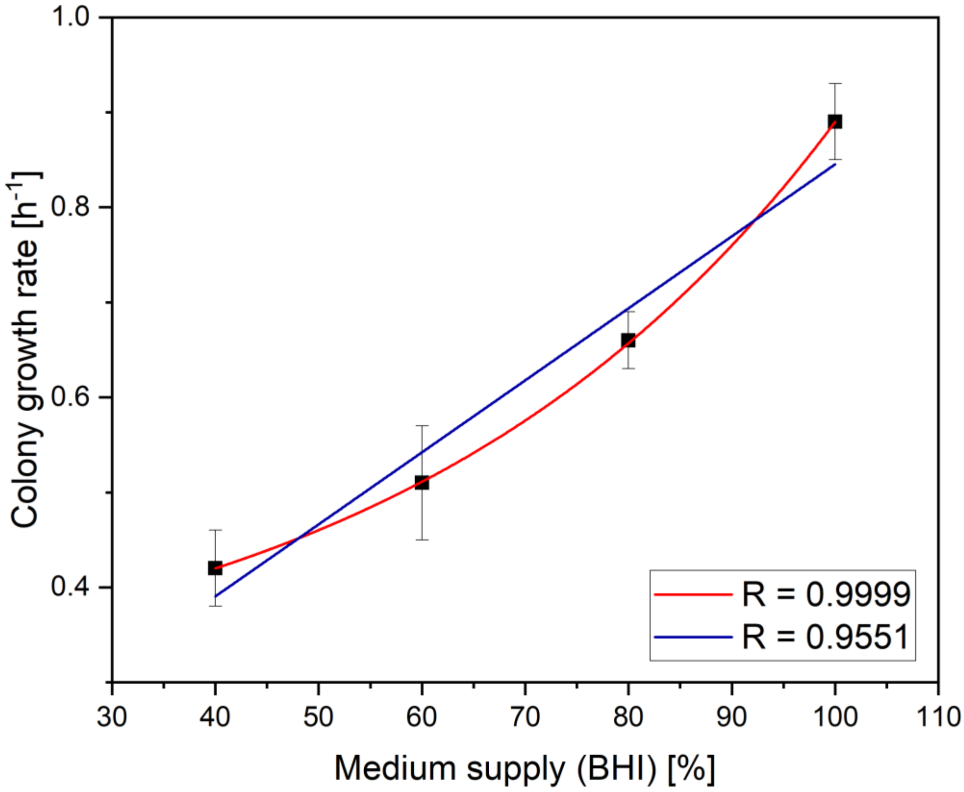


**Fig. S19**: Correlation analysis between the colony growth rate and the overall medium supply by different experimental lifelines.
